# Aberrant mechanosensitive signaling underlies activation of vascular endothelial xanthine oxidoreductase that promotes aortic aneurysm formation in Marfan syndrome

**DOI:** 10.1101/2022.01.30.478356

**Authors:** Hiroki Yagi, Hiroshi Akazawa, Qing Liu, Kimiko Yamamoto, Kan Nawata, Akiko Saga-Kamo, Masahiko Umei, Hiroshi Kadowaki, Ryo Matsuoka, Akito Shindo, Haruhiro Toko, Norifumi Takeda, Masahiko Ando, Haruo Yamauchi, Norihiko Takeda, Mehdi A. Fini, Minoru Ono, Issei Komuro

## Abstract

Marfan syndrome (MFS) is an inherited connective tissue disorder caused by mutations in the *FBN1* gene encoding fibrillin-1, a matrix component of extracellular microfibrils. The main cause of morbidity and mortality in MFS is thoracic aortic aneurysm and dissection, but the underlying mechanisms remain undetermined. We found a significant increase in reactive oxygen species (ROS) generation in ascending aorta of MFS patients and MFS mice harboring the *Fbn1* mutation (C1039G), which was associated with up-regulation of xanthine oxidoreductase (XOR) protein in aortic endothelial cells (ECs). Mechanosensitive signaling involving focal adhesion kinase (FAK)-p38 mitogen-activated protein kinase (MAPK) and early growth response-1 (Egr- 1) was aberrantly activated in ascending aorta of *Fbn1*^C1039G/+^ mice, and mechanical stress on human aortic ECs up-regulated XOR expression through FAK-p38 MAPK activation and Egr-1 up-regulation. Inhibition of XOR function by ECs-specific disruption of *Xdh* gene or by systemic administration of XOR inhibitor febuxostat in *Fbn1*^C1039G/+^ mice suppressed ROS generation, FAK-p38 MAPK activation, and Egr-1 up-regulation, leading to attenuation of aortic aneurysm formation. These findings unveil aberrant mechanosensitive signaling in vascular ECs triggering endothelial XOR activation and ROS generation as a culprit underlying the pathogenesis of aortic aneurysm formation in MFS, and highlight a drug repositioning approach using a uric acid lowering drug febuxostat as a potential therapy for MFS.

## INTRODUCTION

Marfan syndrome (MFS) is a heritable autosomal dominant multisystem disorder of connective tissues affecting 1 in 5,000 to 15,000 individuals ^1,2^. Most premature deaths are related to thoracic aortic aneurysm that grows and progresses to catastrophic aortic rupture or dissection without prior symptoms ^1-3^. Since prophylactic aortic surgery can prolong life expectancy, MFS patients are currently managed by healthy lifestyle measures, treatment with β-adrenergic receptor blockers, and routine surveillance imaging ^1,2,4^. MFS is caused by mutations in *FBN1* gene ^5-7^, which encodes fibrillin-1, a major component of microfibrils in extracellular matrix (ECM) ^8^. Fibrillin-1 not only endows connective tissues with the critical properties of elasticity and resilience, but also controls the bioavailability of transforming growth factor-β (TGF-β), a potent cytokine that regulates proliferation, differentiation, ECM modeling, and apoptosis ^9^. Experimental studies using mouse models for MFS indicated that excessive activation of TGF-β signaling due to insufficient function of fibrillin-1 might be the crucial driver of disease progression in thoracic aortic aneurysm, mitral valve prolapse, and pulmonary emphysema ^10,11^. Although pharmacological inhibition of TGF-β signaling by treatment with angiotensin II type 1 (AT_1_) receptor blocker mitigated aortic aneurysm formation in MFS mice ^12-14^, the pathogenic roles of TGF-β signaling remain elusive ^4,15-17^. It is still unclear which TGF-β pathway, canonical or noncanonical, is involved in MFS-associated aortopathy, and recent studies suggest that TGF-β exerts both protective and detrimental effects on aneurysm ^18^. In addition, several randomized clinical trials demonstrated that AT_1_ receptor blocker provided no superior benefit to β blocker in reducing the rate of aortic dilatation ^4,19-21^.

Reactive oxygen species (ROS) induce endothelial dysfunction, leading to vascular diseases including aortic aneurysms through inflammation, activation of matrix metalloproteinase (MMP), apoptosis of vascular smooth muscle cells (VSMCs) and changes in ECM properties ^22^. Several studies suggested the involvement of ROS generation in the development of thoracic aortic aneurysm in MFS ^23-26^, but the precise roles and mechanisms remain undetermined. ROS are generated in vascular wall by NADPH oxidase, xanthine oxidoreductase (XOR), mitochondrial electron transport chain, and uncoupled endothelial nitric oxide synthase (NOS) ^27^. XOR is the rate-limiting enzyme in purine catabolism that catalyzes the oxidation of hypoxanthine to xanthine and xanthine to uric acid, and exists in two interconvertible forms, xanthine oxidase (XO) and xanthine dehydrogenase (XDH) ^28,29^. XO mainly generates ROS such as superoxide anion and hydrogen peroxide by preferentially using oxygen as the electron acceptor, whereas XDH produces NADH using NAD^+^ as the electron acceptor ^28-30^.

In this study, we demonstrated that endothelial XOR triggered ROS generation in aortic wall and thus promoted thoracic aortic aneurysm formation in MFS. Mechanistically, aberrant activation of mechanosensitive signaling up-regulated XOR expression in vascular endothelial cells (ECs). These results provide mechanistic insights into the pathogenic link how *FBN1* mutations lead to MFS-associated aortopathy.

## RESULTS

### Increased ROS production and up-regulation of endothelial XOR in ascending aorta of MFS mice and patients

*Fbn1*^C1039G/+^ mice showed progressive dilatation of aorta at the levels of aortic root, sinotubular junction (STJ), and ascending aorta, which was accompanied by an increase in the phosphorylation levels of Smad2, extracellular signal-regulated kinase (ERK) 1/2, focal adhesion kinase (FAK), and p38 mitogen-activated protein kinase (MAPK) (Supplementary Fig. 1). In MFS patients, aortic root and ascending aorta were significantly dilated, compared with control patients (Supplementary Fig. 2a, b). Aortic wall was thickened with degeneration of elastic fibers, and deposition of proteoglycan and collagen (Supplementary Fig. 2c), and the phosphorylation levels of Smad3, ERK1/2, FAK, and p38 MAPK were significantly increased (Supplementary Fig.2d, e).

We first evaluated *in situ* ROS generation in ascending aorta of MFS mice harboring the *Fbn1*^C1039G/+^ mice ^31^ and MFS patients. Measurement of dihydroethidium (DHE)- derived fluorescence showed that ROS levels were higher in aortic wall of *Fbn1*^C1039G/+^ mice than in that of control *Fbn1*^+/+^ mice (Fig. 1a). We found that XOR protein expression and XO enzymatic activity were significantly increased in ascending aorta of *Fbn1*^C1039G/+^ mice, compared with *Fbn1*^+/+^ mice (Fig. 1b, c). Immunohistochemical analysis revealed that protein expressions of XOR were specifically elevated in aortic ECs of *Fbn1*^C1039G/+^ mice (Fig. 1d). Similarly, DHE-derived fluorescence was increased in ascending aorta of MFS patients, compared with control patients (Fig. 1e). The mRNA levels of *XDH* gene, encoding XOR, and enzymatic activity of XO were higher in ascending aorta of MFS patients than that of control patients (Fig. 1f, g). Immunohistochemical analysis revealed that XOR protein was up-regulated specifically in aortic ECs of MFS patients (Fig. 1h). These results suggested that an increase in endothelial XOR-derived ROS generation might be involved in aortic aneurysm formation in MFS mice and patients.

**Figure 1.**
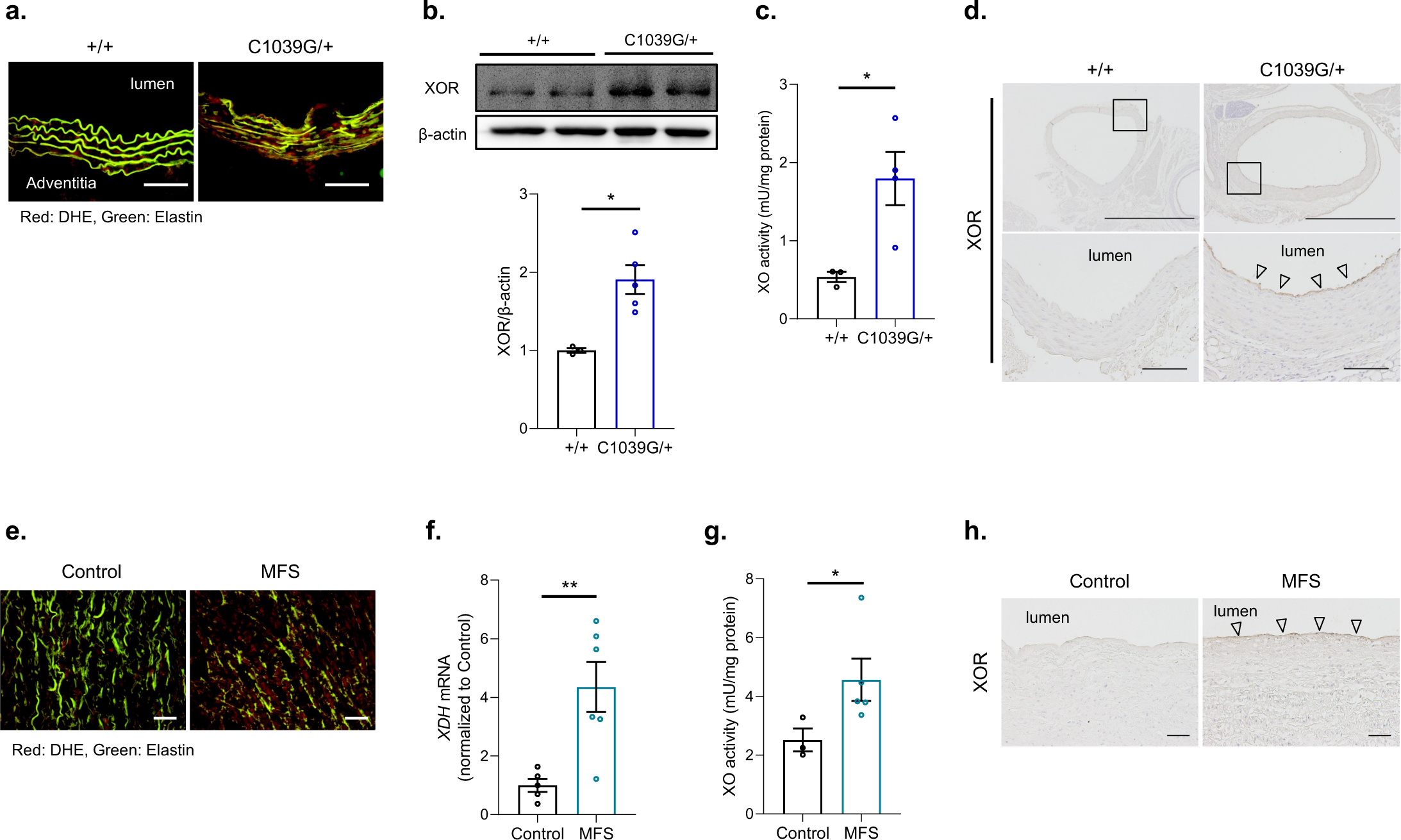
ROS generation and endothelial XOR expression and activity in ascending aorta of MFS mice and patients. **a**. DHE staining of ascending aorta from *Fbn1*^+/+^ and *Fbn1* ^C1039G/+^ mice (16 weeks of age). Merged images with ROS (red) and elastin (green) are shown. Scale bars, 100 μm. **b**. Immunoblot analysis of XOR expression in ascending aorta of *Fbn1*^+/+^ (*n* = 3) and *Fbn1* ^C1039G/+^ mice (*n* = 5) (24 weeks of age). The intensity of each band was quantified by densitometric analysis and corrected for the amount of β-actin protein as an internal control (mean ± SEM). \**P*<0.05, two-tailed Student’s *t*-test. **c**. Measurement of XO enzymatic activity in ascending aorta of *Fbn1*^+/+^ (*n* = 3) and *Fbn1* ^C1039G/+^ mice (*n* = 4) (24 weeks of age). The data are presented as mean ± SEM. \**P*<0.05, two-tailed Student’s *t*-test. **d**. Immunostaining for XOR protein expression in ascending aorta of *Fbn1*^+/+^ and *Fbn1* ^C1039G/+^ mice (24 weeks of age). Scale bars in upper panels, 1 mm; in lower panels, 100 μm. **e**. DHE staining of ascending aorta from MFS patients and control heart transplant recipients. Merged images with ROS (red) and elastin (green) are shown. Scale bars, 50 μm. **f**. The mRNA expressions of *XDH* in ascending aorta of MFS patients (*n* = 6) and control heart transplant recipients (*n* = 5). The data are presented as fold induction over control (mean ± SEM). \**P*<0.05, two-tailed Student’s *t*-test. **g**. Measurement of XO enzymatic activity XO activity in ascending aorta of MFS patients (*n* = 5) and control heart transplant recipients (*n* = 3). The data are presented as mean ± SEM. \**P*<0.05, two-tailed Student’s *t*-test with Welch’s correction. **h**. Immunostaining for XOR protein expression in ascending aorta of MFS patients and control heart transplant recipients. Scale bars, 50 μm.

### Genetic disruption of *Xdh* in endothelial cells suppressed aneurysm formation in *Fbn1*^C1039G/+^ mice

To gain insights into the role of endothelial XOR in the pathogenesis of aortic aneurysm, we inactivated XOR in ECs in *Fbn1*^C1039G/+^ mice. For this purpose, we first generated ECs-specific XOR-knockout mice by crossing *Xdh-*floxed mice with *Tie2* promoter/enhancer-driven *Cre* transgenic mice (Supplementary Fig. 3a), and crossed *Tie2* Cre (+) *Xdh* ^f/f^ mice with *Fbn1*^C1039G/+^ mice. Up-regulation of endothelial XOR protein in *Fbn1*^C1039G/+^ mice was abrogated in *Tie2* Cre (+) *Xdh* ^f/f^ *Fbn1*^C1039G/+^ mice, as revealed by immunostaining and western blot analysis (Supplementary Fig. 3a, b). There were no significant differences in body weight, blood pressure (BP), and pulse rate among *Tie2* Cre (-) *Xdh* ^f/f^ *Fbn1*^+/+^ mice, *Tie2* Cre (-) *Xdh* ^f/f^ *Fbn1*^C1039G/+^ mice, and *Tie2* Cre (+) *Xdh* ^f/f^ *Fbn1*^C1039G/+^ mice (Supplementary Fig. 3c). However, at 16 weeks of age, the enlargement of aortic diameter was significantly suppressed in *Tie2* Cre (+) *Xdh* ^f/f^ *Fbn1*^C1039G/+^ mice, compared with control *Tie2* Cre (-) *Xdh* ^f/f^, *Fbn1*^C1039G/+^ mice (Fig. 2a- c). Histologically, aortic wall thickening with degeneration of elastic fibers and deposition of collagen and proteoglycan were significantly attenuated in ascending aorta of *Tie2* Cre (+) *Xdh* ^f/f^ *Fbn1*^C1039G/+^ mice, compared with control *Tie2* Cre (-) *Xdh* ^f/f^ *Fbn1*^C1039G/+^ mice (Fig. 2d, e). In ascending aorta of *Tie2* Cre (-) *Xdh* ^f/f^ *Fbn1*^C1039G/+^ mice, ROS generation, MMP activation, perivascular infiltration of F4/80-positive macrophages, as well as phosphorylation of Smad2, ERK1/2, FAK, and p38 MAPK were observed, which were also attenuated in *Tie2* Cre (+) *Xdh* ^f/f^ *Fbn1*^C1039G/+^ mice (Fig. 2f-j). These results suggested that up-regulation of XOR in vascular ECs was critically involved in ROS generation in aortic wall and aortic aneurysm formation in *Fbn1*^C1039G/+^ mice.

**Figure 2.**
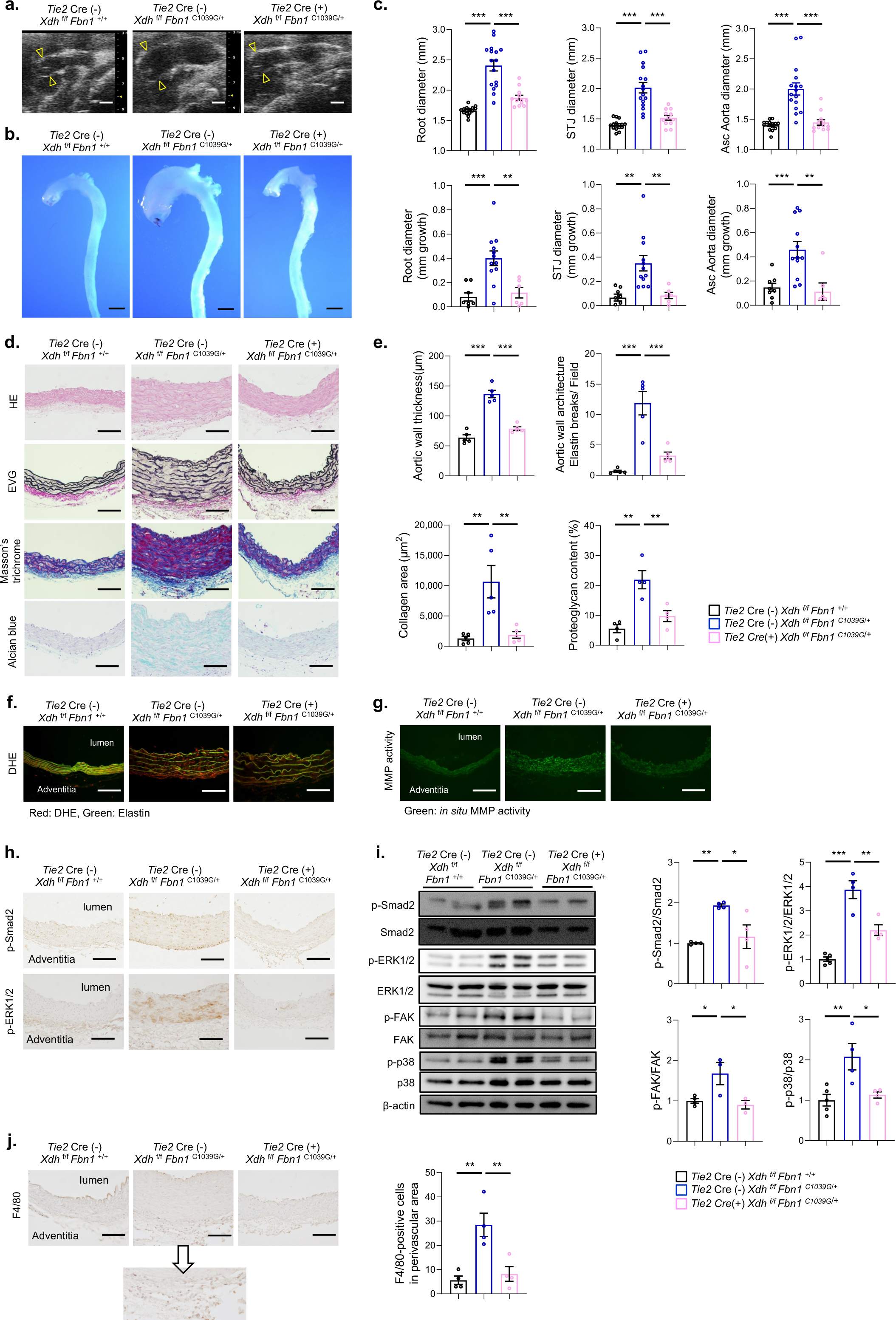
Suppression of aortic aneurysm formation by ECs-specific knockout of *Xdh* in *Fbn1*^C1039G/+^ mice. **a**. Ultrasound images of ascending aorta in *Tie2* Cre (-) *Xdh* ^f/f^ *Fbn1* ^+/+^, *Tie2* Cre (-) *Xdh* ^f/f^ *Fbn1* ^C1039G/+^, and *Tie2* Cre (+) *Xdh* ^f/f^ *Fbn1* ^C1039G/+^ mice (8 weeks of age). Arrowheads indicate aortic root. Scale bars, 1 mm. **b**. Macroscopic appearance of aorta in *Tie2* Cre (-) *Xdh* ^f/f^ *Fbn1* ^+/+^, *Tie2* Cre (-) *Xdh* ^f/f^ *Fbn1* ^C1039G/+^, and *Tie2* Cre (+) *Xdh* ^f/f^ *Fbn1* ^C1039G/+^ mice (16 weeks of age). **c**. Ascending aorta diameters at three different levels (Root, aortic root; STJ, sinotubular junction; Asc Ao, ascending aorta) in 16-week-old *Tie2* Cre (-) *Xdh* ^f/f^ *Fbn1* ^+/+^ (*n* = 15), *Tie2* Cre (-) *Xdh* ^f/f^ *Fbn1* ^C1039G/+^ (n = 16), and *Tie2* Cre (+) *Xdh* ^f/f^ *Fbn1* ^C1039G/+^ (*n* = 12) mice. The data are presented as mean ± SEM. \*\*\**P*<0.001, one-way ANOVA with Tukey’s multiple comparisons test. **d**. Histological analysis with HE staining, elastica van Gieson (EVG) staining, Masson’s trichrome staining, and Alcian blue staining in ascending aorta of *Tie2* Cre (-) *Xdh* ^f/f^ *Fbn1* ^+/+^, *Tie2* Cre (-) *Xdh* ^f/f^ *Fbn1* ^C1039G/+^, and *Tie2* Cre (+) *Xdh* ^f/f^ *Fbn1* ^C1039G/+^ mice (16 weeks of age). Scale bars, 100 μm. **e**. Measurement of aortic wall thickness, aortic wall architecture indicated by the number of breaks of the elastic fiber per field, adventitia collagen area, and proteoglycan content in *Tie2* Cre (-) *Xdh* ^f/f^ *Fbn1* ^+/+^ (*n* = 4 to 5), *Tie2* Cre (-) *Xdh* ^f/f^ *Fbn1* ^C1039G/+^ (*n* = 4 to 5), and *Tie2* Cre (+) *Xdh* ^f/f^ *Fbn1* ^C1039G/+^ mice (*n* = 4 to 5) (16 weeks of age). The data are presented as mean ± SEM. \*\**P*<0.01, \*\*\**P*<0.001, one-way ANOVA with Tukey’s multiple comparisons test. **f**. DHE staining of ascending aorta from *Tie2* Cre (-) *Xdh* ^f/f^ *Fbn1* ^+/+^, *Tie2* Cre (-) *Xdh* ^f/f^ *Fbn1* ^C1039G/+^, and *Tie2* Cre (+) *Xdh* ^f/f^ *Fbn1* ^C1039G/+^ mice (16 weeks of age). Merged images with ROS (red) and elastin (green) are shown. Scale bars, 100 μm. ***g***. *In situ* zymography for gelatinase activity in ascending aorta of *Tie2* Cre (-) *Xdh* ^f/f^ *Fbn1* ^+/+^, *Tie2* Cre (-) *Xdh* ^f/f^ *Fbn1* ^C1039G/+^, and *Tie2* Cre (+) *Xdh* ^f/f^ *Fbn1* ^C1039G/+^ mice (16 weeks of age). Scale bars, 100 μm. **h**. Immunostaining for phosphorylated Smad2 (p-Smad2) and phosphorylated ERK1/2 (p-ERK1/2) in ascending aorta of *Tie2* Cre (-) *Xdh* ^f/f^ *Fbn1* ^+/+^, *Tie2* Cre (-) *Xdh* ^f/f^ *Fbn1* ^C1039G/+^, and *Tie2* Cre (+) *Xdh* ^f/f^ *Fbn1* ^C1039G/+^ mice (16 weeks of age). Scale bars, 100 μm. **i**. Immunoblot analysis of p-Smad2, Smad2, p-ERK1/2, ERK1/2, phosphorylated FAK (p-FAK), FAK, phosphorylated p38 MAPK (p-p38), p38 MAPK (p38), and *β*-actin in ascending aorta of *Tie2* Cre (-) *Xdh* ^f/f^ *Fbn1* ^+/+^, *Tie2* Cre (-) *Xdh* ^f/f^ *Fbn1* ^C1039G/+^, and *Tie2* Cre (+) *Xdh* ^f/f^ *Fbn1* ^C1039G/+^ mice (16 weeks of age). The intensity of each band was quantified by densitometric analysis, and quantitation graphs for the p-Smad2/Smad2 (*n* = 4), p-ERK1/2/ERK1/2 (*n* = 4 to 5), p-FAK/FAK (*n* = 3 to 4), and p-p38/p38 (*n* = 4 to 5) are shown (mean ± SEM). \**P*<0.05, \*\**P*<0.01, \*\*\**P*<0.001, one-way ANOVA with Tukey’s multiple comparisons test. **j**. Immunostaining for F4/80 in ascending aorta of *Tie2* Cre (-) *Xdh* ^f/f^ *Fbn1* ^+/+^, *Tie2* Cre (-) *Xdh* ^f/f^ *Fbn1* ^C1039G/+^, and *Tie2* Cre (+) *Xdh* ^f/f^ *Fbn1* ^C1039G/+^ mice (16 weeks of age). Scale bars, 100 μm. The number of F4/80-positive cells in perivascular area was calculated in each group (*n* = 4). The data are presented as mean ± SEM. \*\**P*<0.01, one-way ANOVA with Tukey’s multiple comparisons test.

### XOR inhibitor febuxostat suppressed aneurysm formation in *Fbn1*^C1039G/+^ mice

We next investigated whether an XOR inhibitor febuxostat [2-(3-cyano-4-isobutoxy- phenyl)-4-methyl-1, 3-thiazole- 5 carboxylic acid] could attenuate progression of aortic root dilatation. *Fbn1*^+/+^ and *Fbn1*^C1039G/+^ mice at 8 weeks of age were administered with vehicle, febuxostat at a lower dose (LD) of 1 mg/kg/day, which is equivalent to the maximum clinical dose ^32,33^, or febuxostat at a higher dose (HD) of 5 mg/kg/day, in their drinking water. *Fbn1*^C1039G/+^ mice treated with either LD or HD febuxostat showed significant improvement in dilatation of aorta at all levels of aortic root, STJ, and ascending aorta, with less elastic fiber fragmentation, and deposition of proteoglycan and collagen, compared with vehicle-treated *Fbn1*^C1039G/+^ mice (Fig. 3a-e). Echocardiographic examination revealed no significant difference in aortic root size among any of the treatment groups for *Fbn1*^C1039G/+^ mice at baseline, but the aortic diameters in *Fbn1*^C1039G/+^ mice was always larger than those in *Fbn1*^+/+^ mice (Fig. 3c). XOR derived-ROS generation and MMP activation, perivascular infiltration of F4/80- positive macrophages, were also attenuated in febuxostat-treated groups (Fig. 3f, g, and j). Febuxostat did not influence BP, pulse rate, body weight, and the levels of serum uric acid and markers for kidney function (Supplementary Fig. 4a, b). Although HD febuxostat inactivated ERK1/2 in *Fbn1*^C1039G/+^ mice more strongly than LD febuxostat, there was no significant difference in inactivation of Smad2, FAK, and p38 MAPK between LD and HD treatment (Fig. 3h, i). Collectively, these results suggested that febuxostat at a dose for clinical use was effective for suppression of aortic aneurysm formation in *Fbn1*^+/+^ mice.

**Figure 3.**
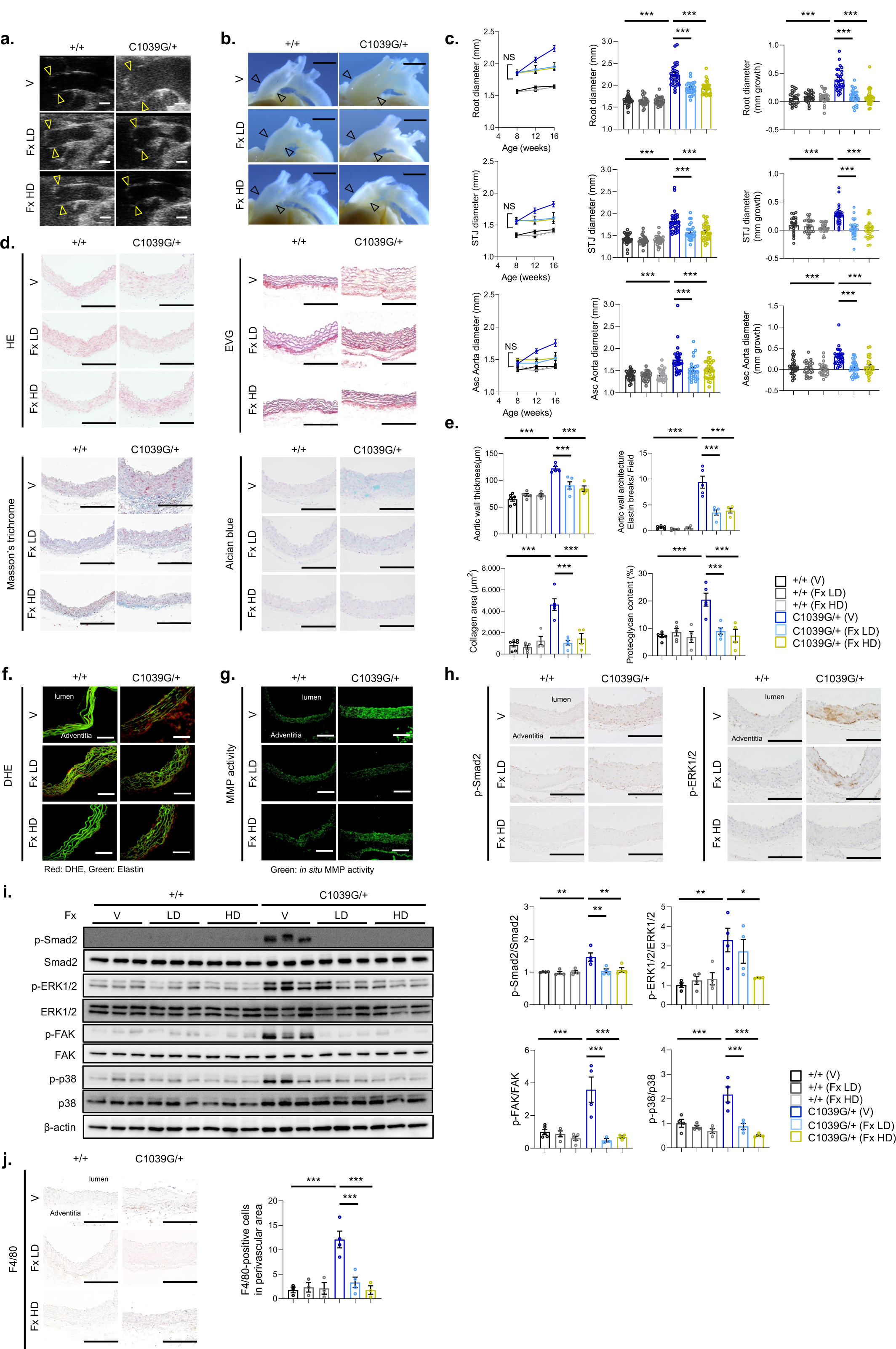
Suppression of aortic aneurysm formation by treatment with febuxostat in *Fbn1*^C1039G/+^ mice. **a**. Ultrasound images of ascending aorta in *Fbn1* ^+/+^ and *Fbn1* ^C1039G/+^ (16 weeks of age) after 8 weeks of treatment with vehicle (V), febuxostat at a lower dose of 1 mg/kg/day (Fx LD), or febuxostat at a higher dose of 5 mg/kg/day (Fx HD). Arrowheads indicate aortic root. Scale bars, 1 mm. **b**. Macroscopic appearance of ascending aorta in *Fbn1* ^+/+^ and *Fbn1* ^C1039G/+^ (16 weeks of age) after 8 weeks of treatment with V, Fx LD, or Fx HD. Arrowheads indicate aortic root. Scale bars, 2 mm. **c**. Ascending aorta diameters at three different levels (Root, aortic root; STJ, sinotubular junction; Asc Ao, ascending aorta) in *Fbn1* ^+/+^ and *Fbn1* ^C1039G/+^ after treatment with V (*Fbn1* ^+/+^; *n* = 25, *Fbn1* ^C1039G/+^; *n* = 28), Fx LD (*Fbn1* ^+/+^; *n* = 25, *Fbn1* ^C1039G/+^; *n* = 26), or Fx HD (*Fbn1* ^+/+^; *n* = 24, *Fbn1* ^C1039G/+^; *n* = 30). Changes in aortic diameters over the 8-week treatment (left panels), aortic diameters at 16 weeks of age (middle panels), and growth of aortic diameters (right panels) are presented as mean ± SEM. \*\*\**P*<0.001, one-way ANOVA with Tukey’s multiple comparisons test. **d**. Histological analysis with HE staining, elastica van Gieson (EVG) staining, Masson’s trichrome staining, and Alcian blue staining in ascending aorta of *Fbn1* ^+/+^ and *Fbn1* ^C1039G/+^ (16 weeks of age) after 8 weeks of treatment with V, Fx LD, or Fx HD. Scale bars, 100 μm. **e**. Measurement of aortic wall thickness, aortic wall architecture indicated by the number of breaks of the elastic fiber per field, adventitia collagen area, and proteoglycan content in *Fbn1* ^+/+^ and *Fbn1* ^C1039G/+^ (16 weeks of age) after 8 weeks of treatment with V, Fx LD, or Fx HD (*n* = 4 to 6). The data are presented as mean ± SEM. \*\*\**P*<0.001, one-way ANOVA with Tukey’s multiple comparisons test. **f**. DHE staining of ascending aorta from *Fbn1* ^+/+^ and *Fbn1* ^C1039G/+^ (16 weeks of age) after 8 weeks of treatment with V, Fx LD, or Fx HD. Merged images with ROS (red) and elastin (green) are shown. Scale bars, 100 μm. ***g***. *In situ* zymography for gelatinase activity in ascending aorta of *Fbn1* ^+/+^ and *Fbn1* ^C1039G/+^ (16 weeks of age) after 8 weeks of treatment with V, Fx LD, or Fx HD. Scale bars, 100 μm. **h**. Immunostaining for phosphorylated Smad2 (p-Smad2) and phosphorylated ERK1/2 (p-ERK1/2) in ascending aorta of *Fbn1* ^+/+^ and *Fbn1* ^C1039G/+^ (16 weeks of age) after 8 weeks of treatment with V, Fx LD, or Fx HD. Scale bars, 100 μm. **i**. Immunoblot analysis of p-Smad2, Smad2, p-ERK1/2, ERK1/2, phosphorylated FAK (p-FAK), FAK, phosphorylated p38 MAPK (p-p38), p38 MAPK (p38), and *β*-actin in ascending aorta of *Fbn1* ^+/+^ and *Fbn1* ^C1039G/+^ (16 weeks of age) after 8 weeks of treatment with V, Fx LD, or Fx HD. The intensity of each band was quantified by densitometric analysis, and quantitation graphs for the p-Smad2/Smad2 (*n* = 4), p-ERK1/2/ERK1/2 (*n* = 4), p-FAK/FAK (*n* = 3 to 5), and p-p38/p38 (*n* = 4) are shown (mean ± SEM). \**P*<0.05, \*\**P*<0.01, \*\*\**P*<0.001, one-way ANOVA with Tukey’s multiple comparisons test. **j**. Immunostaining for F4/80 in ascending aorta of *Tie2* Cre (-) *XOR* ^f/f^ *Fbn1* ^+/+^, *Tie2* Cre (-) *XOR* ^f/f^ *Fbn1* ^C1039G/+^, and *Tie2* Cre (+) *XOR* ^f/f^ *Fbn1* ^C1039G/+^ mice (16 weeks of age). Scale bars, 100 μm. The number of F4/80-positive cells in perivascular area was calculated in each group (*n* = 4). The data are presented as mean ± SEM. \*\**P*<0.01, one-way ANOVA with Tukey’s multiple comparisons test.

### Pressure overload induced phosphorylation of FAK and p38 MAPK more robustly in ascending aorta of *Fbn1*^C1039G/+^ mice than WT mice

We hypothesized that *Fbn1* mutation causes aberrant activation of mechanosensitive signaling such as endothelial XOR activation and ROS generation, leading to aortic aneurysm progression in *Fbn1*^C1039G/+^ mice. Western blot analysis showed a significant increase in the abundance of early growth response-1 (Egr-1), a mechanical stress– responsive transcriptional factor ^34-36^, in ascending aortas of *Fbn1*^C1039G/+^ mice and MFS patients, as compared with *Fbn1*^+/+^ mice and control patients, respectively (Fig. 4a, b). Furthermore, pressure overload by transverse aortic constriction (TAC) induced activation of FAK and p38 MAPK more strongly in the proximal ascending aortas of *Fbn1*^C1039G/+^ mice than in those of *Fbn1*^+/+^ mice at 8 weeks of age (Fig. 4c). These results indicated aberrant activation of mechanosensitive signaling involving the FAK- p38 MAPK pathway in *Fbn1*^C1039G/+^ mice. At 8 weeks of age, the phosphorylation levels of Smad2 and ERK1/2 were comparable, and not further increased after TAC operation in ascending aortas of *Fbn1*^C1039G/+^ and *Fbn1*^+/+^ mice (Fig. 4c).

**Figure 4.**
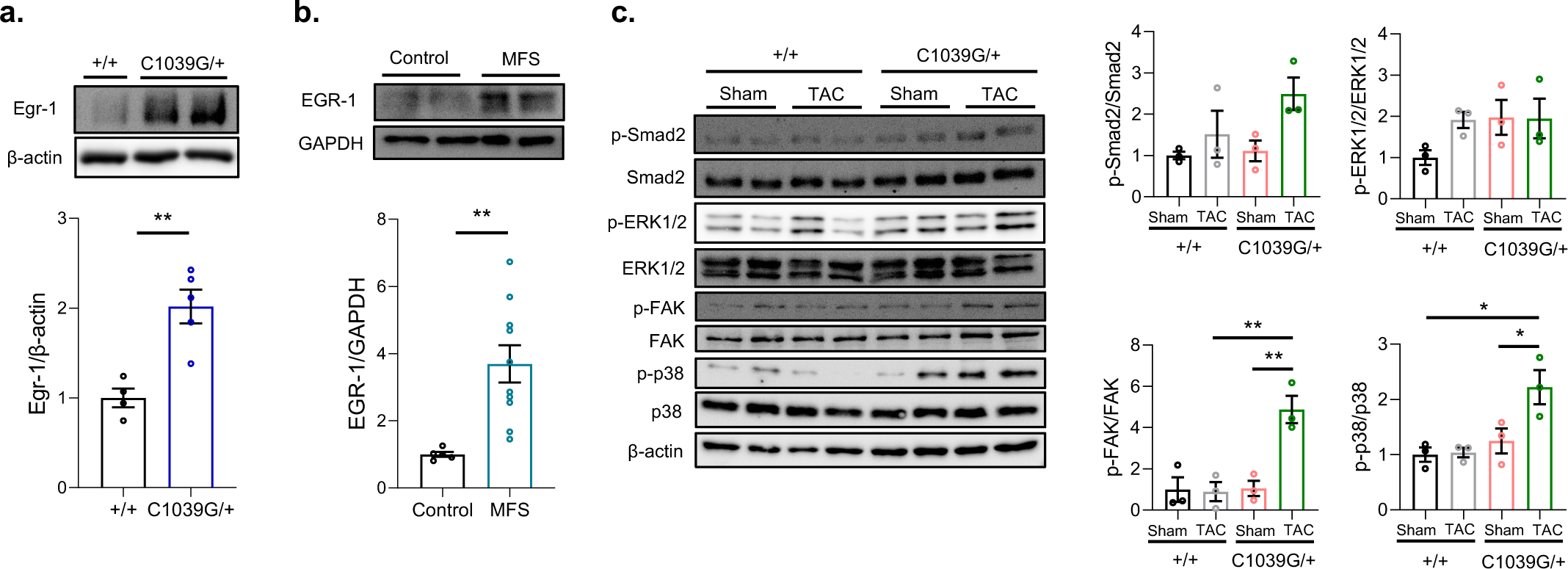
Aberrant activation of mechanosensitive signaling in MFS ascending aorta. **a**. Immunoblot analysis of Egr-1 expression in ascending aorta of *Fbn1*^+/+^ (*n* = 4) and *Fbn1* ^C1039G/+^ mice (*n* = 5) (16 weeks of age). The intensity of each band was quantified by densitometric analysis and corrected for the amount of *β*-actin protein as an internal control (mean ± SEM). \*\**P*<0.01, two-tailed Student’s *t*-test. **b**. Immunoblot analysis of Egr-1 expression in ascending aorta of MFS patients (*n* = 10) and control heart transplant recipients (*n* = 5). The intensity of each band was quantified by densitometric analysis and corrected for the amount of GAPDH protein as an internal control (mean ± SEM). \*\**P*<0.01, two-tailed Student’s *t*-test. **c**. Immunoblot analysis of phosphorylated Smad2 (p-Smad2), Smad2, phosphorylated ERK1/2 (p-ERK1/2), ERK1/2, phosphorylated FAK (p-FAK), FAK, phosphorylated p38 MAPK (p-p38), p38 MAPK (p38), and *β*-actin in ascending aorta of *Fbn1* ^+/+^ (sham; *n* = 3, TAC *n* = 3) and *Fbn1* ^C1039G/+^ (sham; *n* = 3, TAC *n* = 3) (8 weeks of age). The intensity of each band was quantified by densitometric analysis, and quantitation graphs for the p- Smad2/Smad2, p-ERK1/2/ERK1/2, p-FAK/FAK, and p-p38/p38 are shown (mean ± SEM). \**P*<0.05, \*\**P*<0.01, one-way ANOVA with Tukey’s multiple comparisons test.

### XOR was upregulated by mechanical stress through FAK/p38MAPK signaling pathway in endothelial cells

To explore the mechanisms of up-regulation of XOR protein expressions in aortic ECs of *Fbn1*^C1039G/+^ mice, we imposed mechanical stresses on human aortic endothelial cells (HAECs). *XDH* gene was significantly upregulated at 1 h after exposure to hypotonic culture medium (Fig. 5a). Hypotonic stress also induced activation of FAK at 8 min and p38 MAPK at 8 and 15 min, which preceded the up-regulation of *XDH* gene (Fig. 5b). Similarly, cyclic stretch (1 Hz and 10% elongation) significantly increased XOR protein expression at 1 h (Fig. 5c), accompanied by increase in the phosphorylation levels of FAK and p38 MAPK at 8 and 15min (Fig. 5d). To examine the role of FAK and p38 MAPK in up-regulation of XOR, we pretreated HAECs with an FAK inhibitor PF-573228 or a p38 MAPK inhibitor SB-203580. Pretreatment with PF-573228 significantly inhibited hypotonic stress-induced up-regulation of XOR protein, as well as activation of FAK and p38 MAPK in HAECs (Fig. 5e, f). Pretreatment with SB-203580 did not affect the phosphorylation levels of FAK, but significantly inhibited hypotonic stress-induced up- regulation of XOR protein (Fig. 5g, h). These results suggested that up-regulation of XOR was induced by mechanical stress through the FAK/p38 MAPK signaling pathway in HAECs.

**Figure 5.**
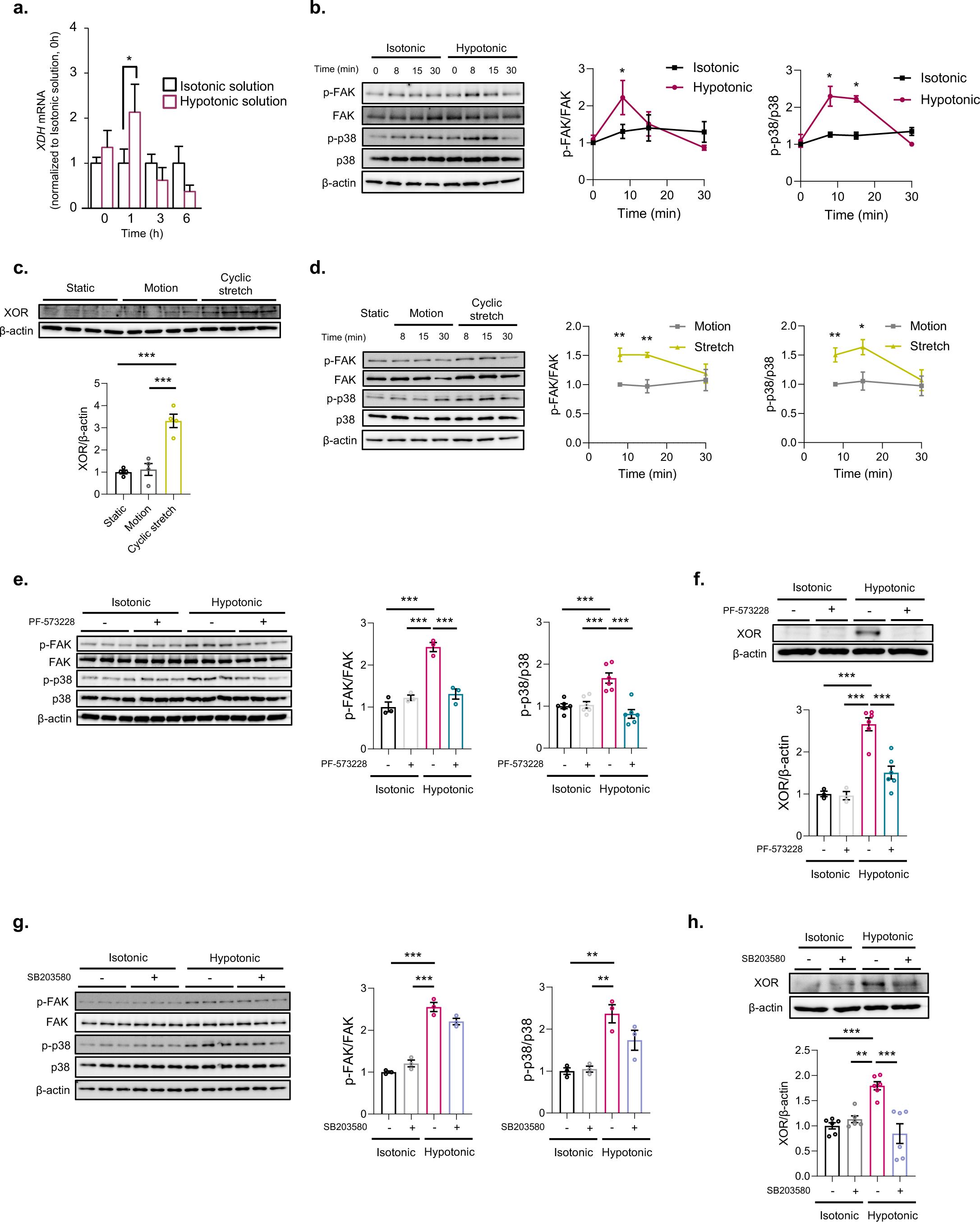
Mechanical stress-induced up-regulation of XOR through FAK/p38MAPK signaling pathway in endothelial cells. **a**. The mRNA expressions of *XDH* in HAECs at 0, 1, 3, and 6 h after exposure to isotonic or hypotonic solution (*n* = 4). The data are presented as fold induction over Isotonic solution at 0 h (mean ± SEM). \**P*<0.05, two-tailed Student’s *t*-test. **b**. Immunoblot analysis of phosphorylated FAK (p-FAK), FAK, phosphorylated p38 MAPK (p-p38), p38 MAPK (p38), and β-actin in HAECs at 0, 8, 15, and 30 min after exposure to isotonic or hypotonic solution (*n* = 3). The intensity of each band was quantified by densitometric analysis, and quantitation graphs for the p-FAK/FAK, and p- p38/p38 are shown (mean ± SEM). \**P*<0.05, two-tailed Student’s *t*-test. **c**. Immunoblot analysis of XOR expression in HAECs after 1 h under static, motion, or cyclic stretch (1 Hz, 10% elongation) conditions (*n* = 4). The intensity of each band was quantified by densitometric analysis and corrected for the amount of β-actin protein as an internal control (mean ± SEM). \*\*\**P*<0.001, one-way ANOVA with Tukey’s multiple comparisons test. **d**. Immunoblot analysis of phosphorylated FAK (p-FAK), FAK, phosphorylated p38 MAPK (p-p38), p38 MAPK (p38), and β-actin in HAECs after 8, 15, and 30 min under motion or cyclic stretch (1 Hz, 10% elongation) conditions (*n* = 4). The intensity of each band was quantified by densitometric analysis, and quantitation graphs for the p- FAK/FAK, and p-p38/p38 are shown (mean ± SEM). \**P*<0.05, \*\**P*<0.01, two-tailed Student’s *t*-test. **e**. Immunoblot analysis of phosphorylated FAK (p-FAK), FAK, phosphorylated p38 MAPK (p-p38), p38 MAPK (p38), and β-actin in HAECs in HAECs at 8 min after exposure to isotonic or hypotonic solution with or without pretreatment with an FAK inhibitor PF-573228 (*n* = 3 to 6). The intensity of each band was quantified by densitometric analysis, and quantitation graphs for the p-FAK/FAK, and p-p38/p38 are shown (mean ± SEM). \*\*\**P*<0.001, one-way ANOVA with Tukey’s multiple comparisons test. **f**. Immunoblot analysis of XOR expression in HAECs at 1 h after exposure to isotonic or hypotonic solution with or without pretreatment with PF-573228 (*n* = 3 to 6). The intensity of each band was quantified by densitometric analysis and corrected for the amount of β-actin protein as an internal control (mean ± SEM). \*\*\**P*<0.001, one-way ANOVA with Tukey’s multiple comparisons test. **g**. Immunoblot analysis of phosphorylated FAK (p-FAK), FAK, phosphorylated p38 MAPK (p-p38), p38 MAPK (p38), and β-actin in HAECs at 8 min after exposure to isotonic or hypotonic solution with or without pretreatment with a p38 MAPK inhibitor SB-203580 (*n* = 3). The intensity of each band was quantified by densitometric analysis, and quantitation graphs for the p-FAK/FAK, and p-p38/p38 are shown (mean ± SEM). \*\**P*<0.01, \*\*\**P*<0.001, one-way ANOVA with Tukey’s multiple comparisons test. **h**. Immunoblot analysis of XOR expression in HAECs at 1 h after exposure to isotonic or hypotonic solution with or without pretreatment with PF-573228 (*n* = 6). The intensity of each band was quantified by densitometric analysis and corrected for the amount of β- actin protein as an internal control (mean ± SEM). \*\**P*<0.01, \*\*\**P*<0.001, one-way ANOVA with Tukey’s multiple comparisons test.

### Egr-1 mediated mechanical stress-induced up-regulation of XOR in MFS

Finally, we investigated how the FAK/p38 MAPK signaling pathway led to up-regulation of XOR in response to mechanical stress. We confirmed that the expression levels of Egr-1 protein were significantly increased by hypotonic stress and cyclic stretch in HAECs (Fig. 6a, b). Pharmacological inhibition of p38 MAPK by treatment with SB- 203580 abrogated hypotonic stress-induced up-regulation of Egr-1 protein in HAECs, indicating that Egr-1 was a mechanical stress-responsive target downstream of p38 MAPK (Fig. 6c). To examine the role of Egr-1 in mechanical stress-induced up- regulation of XOR protein, we knocked down the *EGR1* gene by siRNA in HAECs. siRNA knockdown of Egr-1 abrogated up-regulation of XOR protein, as well as Egr-1 protein, in response to hypotonic stress and cyclic stretch (Fig. 6d-g), indicating that Egr- 1 was critically involved in up-regulation of XOR protein in response to mechanical stress-induced activation of the FAK/p38 MAPK signaling pathway in HAECs.

**Figure 6.**
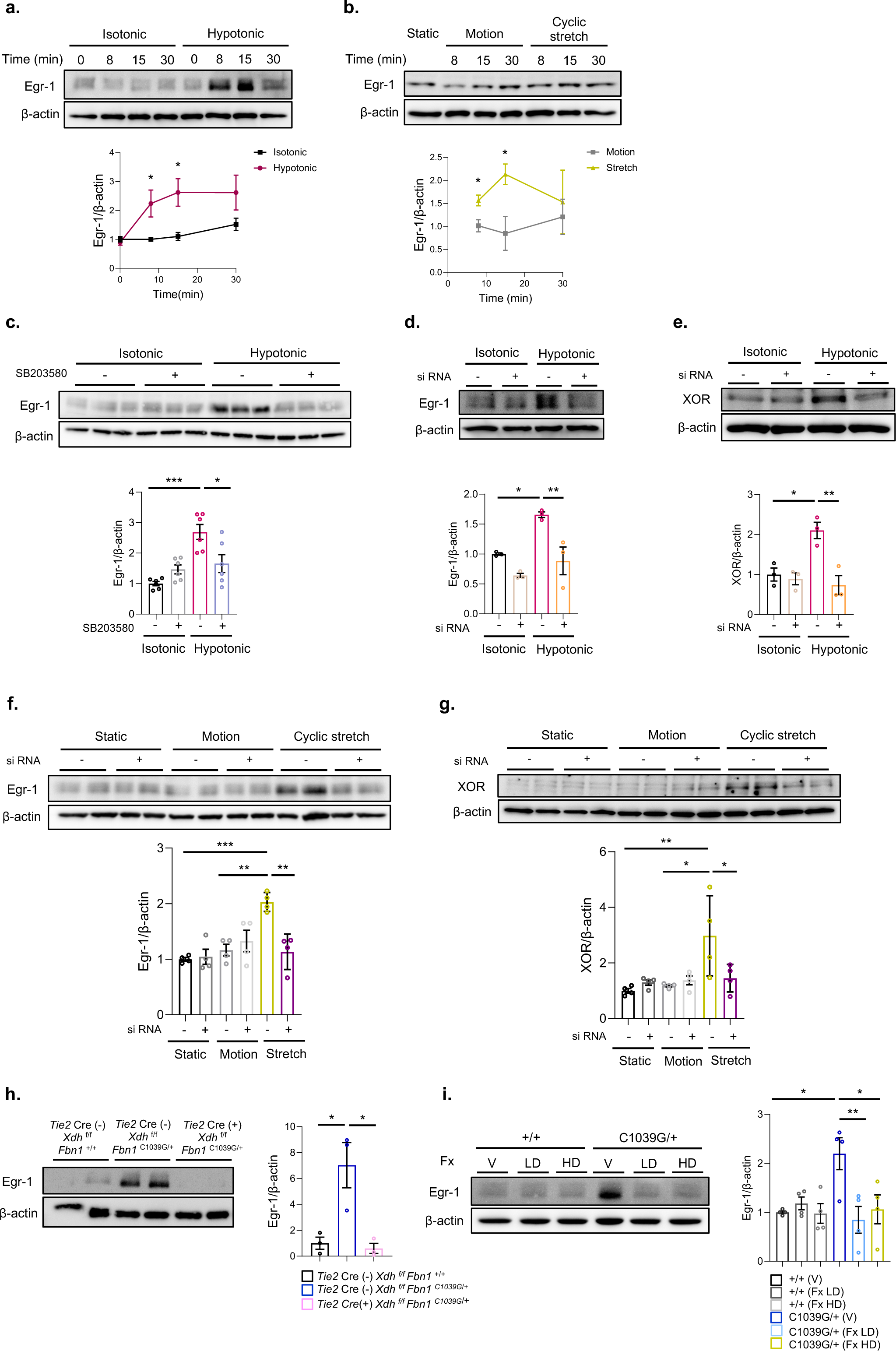
Involvement of Egr-1 in up-regulation of XOR in response to mechanical stress-induced activation of FAK/p38 MAPK signaling pathway in endothelial cells. **a**. Immunoblot analysis of Egr-1 expression in HAECs at 0, 8, 15, and 30 min after exposure to isotonic or hypotonic solution (*n* = 5). The intensity of each band was quantified by densitometric analysis and corrected for the amount of β-actin protein as an internal control (mean ± SEM). \**P*<0.05, two-tailed Student’s *t*-test. **b**. Immunoblot analysis of Egr-1 expression in HAECs after 8, 15, and 30 min under motion or cyclic stretch (1 Hz, 10% elongation) conditions (*n* = 3). The intensity of each band was quantified by densitometric analysis and corrected for the amount of β- actin protein as an internal control (mean ± SEM). \**P*<0.05, two-tailed Student’s *t*-test. **c**. Immunoblot analysis of Egr-1 expression in HAECs at 8 min after exposure to isotonic or hypotonic solution with or without pretreatment with a p38 MAPK inhibitor SB- 203580 (*n* = 6). The intensity of each band was quantified by densitometric analysis and corrected for the amount of β-actin protein as an internal control (mean ± SEM). \**P*<0.05, \*\*\**P*<0.001, one-way ANOVA with Tukey’s multiple comparisons test. **d**. Immunoblot analysis of Egr-1 expression in HAECs at 15 min after exposure to isotonic or hypotonic solution with or without siRNA-mediated knockdown of Egr-1 (*n* = 3). The intensity of each band was quantified by densitometric analysis and corrected for the amount of β-actin protein as an internal control (mean ± SEM). \**P*<0.05, \*\**P*<0.01, one-way ANOVA with Tukey’s multiple comparisons test. **e**. Immunoblot analysis of XOR expression in HAECs at 1 h after exposure to isotonic or hypotonic solution with or without siRNA-mediated knockdown of Egr-1 (*n* = 3). The intensity of each band was quantified by densitometric analysis and corrected for the amount of β-actin protein as an internal control (mean ± SEM). \**P*<0.05, \*\**P*<0.01, one-way ANOVA with Tukey’s multiple comparisons test. **f**. Immunoblot analysis of Egr-1 expression in HAECs after 15 min under static, motion, or cyclic stretch (1 Hz, 10% elongation) conditions with or without siRNA-mediated knockdown of Egr-1 (*n* = 4). The intensity of each band was quantified by densitometric analysis and corrected for the amount of β-actin protein as an internal control (mean ± SEM). \*\**P*<0.01, \*\*\**P*<0.001, one-way ANOVA with Tukey’s multiple comparisons test. **g**. Immunoblot analysis of XOR expression in HAECs after 1 hr under static, motion, or cyclic stretch (1 Hz, 10% elongation) conditions with or without siRNA-mediated knockdown of Egr-1 (*n* = 4). The intensity of each band was quantified by densitometric analysis and corrected for the amount of β-actin protein as an internal control (mean ± SEM). \**P*<0.05, \*\**P*<0.01, one-way ANOVA with Tukey’s multiple comparisons test. **h**. Immunoblot analysis of Egr-1 expression in ascending aorta of *Tie2* Cre (-) *Xdh* ^f/f^ *Fbn1*^+/+^, *Tie2* Cre (-) *Xdh* ^f/f^ *Fbn1* ^C1039G/+^, and *Tie2* Cre (+) *Xdh* ^f/f^ *Fbn1* ^C1039G/+^ mice (*n* = 3) (16 weeks of age). The intensity of each band was quantified by densitometric analysis and corrected for the amount of β-actin protein as an internal control (mean ± SEM). \**P*<0.05, one-way ANOVA with Tukey’s multiple comparisons test. **i**. Immunoblot analysis of Egr-1 expression in ascending aorta of *Fbn1* ^+/+^ and *Fbn1* ^C1039G/+^ (16 weeks of age) after 8 weeks of treatment with V, Fx LD, or Fx HD (*n* = 4). The intensity of each band was quantified by densitometric analysis and corrected for the amount of β-actin protein as an internal control (mean ± SEM). \**P*<0.05, \*\**P*<0.01, one-way ANOVA with Tukey’s multiple comparisons test.

The expression levels of Egr-1 protein were significantly increased in ascending aorta of *Fbn1*^C1039G/+^ mice, as compared with *Fbn1*^+/+^ mice (Fig. 4a). Interestingly, inactivation of XOR either by ECs-specific knockout or systemic administration of febuxostat abrogated up-regulation of Egr-1 protein in *Fbn1*^C1039G/+^ mice (Fig. 6h, i), indicating that inhibition of *in vivo* function of XOR prevented aortic aneurysm formation, and thus decreased wall stress in the ascending aorta. Collectively, these results suggested that mechanical stress activated the FAK/p38 MAPK signaling pathway and up-regulated Egr-1 and XOR proteins in vascular ECs, and that a vicious cycle of mechanical stress and endothelial up-regulation of XOR protein underlie the pathogenesis of aortic aneurysm in MFS.

## DISCUSSION

In this study, we demonstrated that aberrant activation of mechanosensitive signaling in vascular ECs triggered endothelial XOR activation and ROS generation, leading to progression of aneurysm formation in ascending aorta of MFS. Up-regulation and activation of endothelial XOR protein and higher DHE fluorescence were observed in ascending aorta of both MFS mice and patients, and genetic disruption of endothelial XOR or pharmacological inhibition of XOR by febuxostat attenuated aortic aneurysm formation in MFS mice.

ROS play a central role in the development of not only atherosclerotic abdominal aortic aneurysm ^37^, but also genetically triggered thoracic aortic aneurysm ^22,38^. Increased levels of ROS production in aortic wall lead to a wide range of molecular and cellular alterations including activation of MMPs and induction of VSMCs apoptosis and inflammation, and thereby contributes to aortic aneurysm formation ^22,37^. It was reported that protein carbonyl content in plasma of MFS patients was significantly elevated and showed a positive correlation with the clinical score, while total antioxidant capacity was reduced, indicating the involvement of oxidative stress-mediated damage in MFS ^24^. Oxidative stress was accumulated in ascending aorta of MFS mice and patients, as revealed by an increase in the levels of oxidative stress markers such as 8-isoprostane and nitrotyrosine residues ^23,25^. Among the ROS-producing enzymes, the expression levels of NADPH oxidase, inducible NOS (NOS2), and XOR were increased in MFS aorta ^23,25^, and genetic disruption of *Nos2* or *Nox4* ameliorated aortic pathology in *Fbn1*^C1039G/+^ mice ^25,39^. In our study, DHE fluorescence was distributed not only in intima but also in media of MFS aorta (Fig. 1a, e), and was uniformly diminished by genetic disruption of endothelial XOR (Fig. 2f), as well as by treatment with febuxostat (Fig. 3f). We suppose that XOR-derived ROS in intimal ECs may amplify ROS generation by propagating signals to the neighboring cells, and consequently activating ROS-producing enzymes such as NADPH oxidase and inducible NOS in medial layer of the aorta in MFS. Indeed, flow-mediated dilatation, a noninvasive measurement of endothelial function, indicated endothelial dysfunction in MFS patients ^40,41^, and the development of aortic aneurysm in *Fbn-1* hypomorphic *Fbn1*^mgR/mgR^ mice was mitigated by disruption of *Agtr1a* gene encoding angiotensin II type 1a receptor in intimal ECs, but not in medial smooth muscle cells (SMCs) ^42^, These data indicate that endothelial and smooth muscle cell interactions underlie the pathophysiology of aortic aneurysm formation in MFS. Importantly, ROS modulate TGF-β signaling at the levels of gene expression and latent TGF-β activation, and TGF-β reciprocally increases ROS by stimulating mitochondrial ROS production, inducing NADPH oxidases, and suppressing antioxidant enzyme activities ^43^. In our study, the phosphorylation levels of Smad2 and ERK1/2 in aortic wall of MFS mice were uniformly reduced by genetic disruption of endothelial XOR (Fig. 2h, i) or febuxostat treatment (Fig. 3h, i), suggesting a role of the vicious cycle between ROS and TGF-β in amplifying ROS generation throughout the aortic wall (Supplementary Fig. 5).

The enzymatic activity of XOR is determined by a combination of transcriptional regulation and post-translational modification ^28,29^. The XOR expression is up- regulated by a variety of factor such as inflammatory cytokines, lipopolysaccharide, growth factors, glucocorticoids, and hypoxia ^28,30^. It was reported that XOR-dependent ROS production was also increased by oscillatory shear stress in bovine aortic endothelial cells, which was associated with an elevated ratio of XO to XDH ^44^. In this study, we showed that XOR expression was up-regulated by hypotonic stress or cyclic stretch through the FAK/p38 MAPK signaling pathway in HAECs (Fig. 5). In addition, a mechanical stress–responsive transcriptional factor Egr-1 mediated up-regulation of XOR protein in response to mechanical stress-induced activation of the FAK/p38 MAPK signaling pathway (Fig. 6). Mutations in genes encoding ECM proteins can impair structural integrity and matrix-cell interactions in vessel wall, and thereby alter responses to mechanical stress ^36^. A vascular model using human induced pluripotent stem cells (hiPSC) recapitulated the cellular pathologies observed in MFS, and cyclic stretch induced higher density of stress fibers and focal adhesions and higher activation of p38 MAPK in MFS patient-specific hiPSC-derived SMCs than in controls ^45^. Aberrant activation of mechanosensitive signaling was also confirmed in our study, demonstrating an increase in Egr-1expression in ascending aorta of MFS mice and patients (Fig. 4a, b) and an increase in phosphorylation levels of FAK and p38 MAPK after TAC operation in ascending aorta of MFS mice (Fig. 4c). Interestingly, activation of the mechaosensitive signaling involving FAK, p38 MAPK, and Egr-1 in MFS mice was suppressed by genetic disruption of endothelial XOR (Fig. 2i and 6h) or febuxostat treatment (Fig. 3i and 6i). According to Laplace’s law, an increase in the inner radius of vessels proportionally increases circumferential wall stress ^36,46^. We assume that inhibition of endothelial XOR attenuates progression of aortic aneurysm formation, and prevention of aortic dilatation further decreases wall stress and aberrant mechanosensitive signaling, breaking the vicious cycle between abnormal mechanotransduction and aortic aneurysm formation (Supplementary Fig. 5).

Recent research effort advanced our understanding of the pathogenesis of MFS aortopahy, but there has been no progress in drug therapy for a long time ^4,18,20^. Treatment with β-blocker has been recommended to reduce the rate of aortic dilatation in MFS patients ^47,48^. Although an AT_1_ receptor irbesartan was reported to reduce aortic dilatation compared with placebo ^49^, superior therapeutic effect of AT_1_ receptor blocker to β-adrenergic receptor blocker has not been conclusively proven by large randomized clinical trials ^4,18,20,21^. Preclinical studies using *Fbn1*^C1039G/+^ or *Fbn1*^mgR/mgR^ mice as a disease model have identified several potential targets for new drug therapy ^4^. Based on the present study, we propose potential use of febuxostat for the prevention of aortic dilatation in MFS patients. Febuxostat is approved for the management of hyperuricemia. Although a recent clinical trial raised concerns regarding an increase in the risk of cardiovascular mortality in patients with cardiovascular comorbidities who were treated with febuxostat ^50^, a recent prospective and randomized trial in patients with gout showed that febuxostat was non-inferior to allopurinol therapy with respect to the primary cardiovascular endpoint, and its long-term use was not associated with an increased risk of death or serious adverse events, compared with allopurinol ^51^. In our study, treatment with febuxostat at a clinical dose provided preventive effects on aortic aneurysm formation in *Fbn1*^C1039G/+^ mice, and importantly, XOR protein was also up- regulated in aortic ECs of MFS patients (Fig. 1f, h).

In conclusion, aberrant activation mechanosensitive signaling in vascular ECs triggered endothelial XOR activation and ROS generation, which contributes to the pathogenesis of aortic aneurysm formation in MFS. Our results highlight a drug repositioning approach using a uric acid lowering drug febuxostat as a potential therapy for aortic aneurysm in MFS patients.

## METHODS

### Animal experiments

All animal experiments were approved by the Animal Care Committee of The University of Tokyo, and carried out in accordance with the animal experiment guidelines. Mice are housed in groups of 3-4 animals per cage with a 12 h light-dark cycle at constant room temperature of 22 ± 1 °C with a humidity of 50 ± 15 %. Mice are fed a standard chow (CE-2; Clea Japan Inc.) and water *ad libitum* during the experimental period. *Fbn1* ^C1039G/+^ and *Tie2*-*Cre* mice were obtained from the Jackson Laboratory ^31,52^, and maintained on a C57BL/6J background. *Xdh* ^f/f^ mice were developed by Dr. Mehdi Fini and described previously ^53,54^. Febuxostat was synthesized in Teijin Pharma Limited, and administered via drinking water (1 mg/kg/day or 5 mg/kg/day) ^32,33^. We imposed pressure overload on 8-week-old mice by TAC operation, as described previously ^55^. Transverse aorta was constricted with a 7-0 polypropylene suture by ligating the aorta with splinting a 25-gauge blunted needle, which was removed after the ligation. For evaluation of aortic dimensions as well as cardiac dimensions and contractility, transthoracic echocardiography was performed using a Vevo 2100 system with a 30-MHz probe (FUJIFILM Visualsonics, Inc.). Mice were anesthetized with 2-3% isoflurane inhalation during echocardiographic assessment. A two-dimensional short-axis view of aorta was obtained for measurement of the aortic diameters at the level of aortic root, ST junction, and ascending aorta. The blood pressures and pulse rates were measured in conscious mice noninvasively using the tail-cuff system (MK-2000ST NP-NIBP Monitor; Muromachi Kikai Co., Ltd.). Serum concentrations of uric acid and markers for kidney function (blood urea nitrogen and creatinine) were measured in the laboratory of SRL, Inc.

### Human aortic tissue specimens

This study was approved by the Research Ethics Committee, Graduate School of Medicine and Faculty of Medicine, The University of Tokyo (Reference No. 2233-7). Ascending aortic aneurysm samples were collected from MFS patients undergoing elective or emergent aortic root surgery (Supplementary Table). All patients fulfilled MFS diagnostic criteria according to the revised Ghent nosology ^56^. Control aortic tissues were obtained from heart transplant recipients undergoing transplantation after obtaining written informed consent (Supplementary Table). To evaluate histological and immunohistochemical analysis, aortic tissues were either fixed in 10% formalin (Sakura Finetek Japan) and embedded in paraffin, or immediately embedded in Tissue- Tek OCT compound (Miles Laboratories, Inc.). For western blot and real-time quantita- tive RT–PCR analysis, aortic tissues were immediately immersed in liquid nitrogen and preserved at -80°C in deep freezer.

### Western blot analysis

Aortic tissues were harvested from mice, and perivascular adipose tissue was thoroughly removed. Aortas were divided into ascending thoracic (from the aortic root to immediately past the left subclavian artery) and descending thoracic aortas. Human and murine aortas were crushed by cryo-press, and dissolved in RIPA lysis buffer (50 mM Tris-HCl, pH 8.0, 150 mM NaCl, 1% NP-40, 1% SDS, 0.5% Na-deoxycholate, 10 mM Okadaic acid) containing protease inhibitor cocktail (cOmplete ULTRA Tablets Mini EASYpack; Roche Diagnostics). Protein concentration was measured by BCA Protein Assay Kit (Thermo Fisher Scientific). The lysates were fractionated with SDS– PAGE, transferred to polyvinylidene difluoride membrane (Merck Millipore). The blotted membranes were incubated with the following primary antibodies: rabbit monoclonal anti-phosphorylated Smad2 (Ser465/467) antibody (Cell Signaling Technology, #3108), rabbit monoclonal anti-Smad2 antibody (Cell Signaling Technology, #5339), rabbit monoclonal anti-phosphorylated Smad3 (Ser423/425) antibody (Abcam, #ab52903), rabbit monoclonal anti-Smad3 antibody (Cell Signaling Technology, #9523), rabbit polyclonal anti-phosphorylated ERK1/2 (Thr202/Tyr204) antibody (Cell Signaling Technology, #9101), rabbit polyclonal anti-ERK1/2 antibody (Cell Signaling Technology, #9102), rabbit monoclonal anti-phosphorylated p38 MAPK antibody (Thr180/Tyr182) (Cell Signaling Technology, #4511), rabbit polyclonal anti-p38 MAPK antibody (Cell Signaling Technology, #9212), mouse monoclonal anti-phosphorylated FAK antibody (Tyr397) (Abcam, #ab82198), rabbit polyclonal anti-FAK antibody (Cell Signaling Technology, #3285), mouse monoclonal anti-XO antibody (Santa Cruz Biotechnology, Inc., #sc-398548), rabbit polyclonal anti-XO antibody (Abcam, #ab176165), rabbit monoclonal anti-EGR1 antibody (Cell Signaling Technology, #4154), mouse monoclonal anti-β-actin antibody (Sigma-Aldrich, #A2228), and rabbit monoclonal anti-GAPDH antibody (Cell Signaling Technology, #2118). Membranes were then incubated with horseradish peroxidase-conjugated anti-mouse (Jackson Immuno Research Laboratories, Inc., #115-035-146) or anti-rabbit IgG antibody (Jackson Immuno Research Laboratories, Inc., # 111-035-003). Immunoreactive signals were detected with ECL Prime Western Blotting Detection Reagent (GE Healthcare Biosciences), and visualized with Lumino Graph I (ATTO Corp.). Images were converted to gray scale, and the integrated density per image area of interest was quantified using an NIH Image J software (NIH, Research Branch).

### Real-time quantitative RT-PCR

Total RNA was isolated from cultured cells or tissues by using the Trizol Reagent (Thermo Fisher Scientific). Single-stranded cDNA was reverse transcribed using Prime Script RT Master Mix (Perfect Real Time) (Takara Bio Inc.), and used for subsequent real-time PCR reactions. Gene expression was assessed using SYBR green- based quantification (Roche Diagnostics). PCRs were carried out in a LightCycler 480 system (Roche Diagnostics) with initial denaturation for 3 min at 95°C and then 39 cycles of 10 s at 95°C and 30 s at 55°C. The expression level of a gene was standardized to that of *GAPDH* using the ΔCt method. The PCR primers used were as follows: *XDH* (5’-TGCCCAAAAGACAGAGGTGT-3’, and 5’-CAGTGATGATGTTCCCTCCA-3’), *EGR1* (5’-AGCCCTACGAGCACCTGAC-3’, and 5’-GGTTTGGCTGGGGTAACTG- 3’), *GAPDH* (5’-CCCCGGTTTCTATAAATTGAGC-3’, and 5’- CACCTTCCCCATGGTGTCT-3’)

### Histological and immunohistochemical analysis

For histological analysis, the ascending aorta was excised, fixed immediately in 10% neutralized formalin (Sakura Finetek Japan Co., Ltd.), and embedded in paraffin. Serial sections at 5μm were stained with hematoxylin-eosin (HE) for morphological analysis, elastica van Gieson (EVG) for detection of elastic fibers, Masson’s trichrome for detection of collagen fibers, and Alcian blue for staining proteoglycan. Aortic wall architecture was assessed using a scale of 1 (no break in the elastic fiber) through 4 (diffuse disruption of the elastic fiber) ^12^. Breaks of elastic fiber and deposition of proteoglycan and collagen were assessed at 4-5 different representative locations and averaged by 2 observers blinded to the genotype and treatment for each mouse.

For immunohistochemical analysis, deparaffinized sections were rehydrated, and boiled to retrieve antigens (10 mM citrate buffer, pH 6.0 for staining of phosphorylated Smad2, ERK1/2, p38 MAPK and FAK; 0.1% trypsin solution for staining of F4/80). After blocking with 10% goat or rabbit serum in PBS and with Avidin/Biotin Blocking Kit (Vector Laboratories, Inc.), sections were incubated with the following primary antibodies: rabbit polyclonal anti-XO antibody (Santa Cruz Biotechnology, Inc., #sc- 20991), rabbit monoclonal anti-phosphorylated Smad2 antibody (Ser465/467) (Cell Signaling Technology, #3108), rabbit polyclonal anti-phosphorylated ERK1/2 antibody (Thr202/Tyr 204) (Cell Signaling Technology, #9101), rabbit monoclonal anti- phosphorylated p38 MAPK antibody (Thr180/Tyr182) (Cell Signaling Technology, #4511), rabbit polyclonal anti-phosphorylated FAK (Tyr397) antibody (Cell Signaling Technology, #3283), and rat monoclonal anti-F4/80 antibody (Bio-Rad Laboratories, Inc., MCA497GA). Vectastain ABC kit (Vector Laboratories, Inc.) and DAB Peroxidase Substrate Kit (Vector Laboratories, Inc.) were used to detect the primary antibodies. The sections were counterstained with hematoxylin and mounted in Mount-Quick (Daido Sangyo Co., Ltd.).

### Measurement of XO activity

We measured XO activity with a fluorometric pterin-based assay, as previously described ^32,57^. Aortic tissues were homogenized in 50 mM potassium phosphate buffer (50 mM KH_2_PO_4_, 50 mM Na_2_HPO_4,_ pH 7.4) containing 1 mM EDTA and protease inhibitor cocktail (cOmplete ULTRA Tablets Mini EASYpack; Roche Diagnostics). Protein concentration was measured by BCA Protein Assay Kit (Thermo Fisher Scientific). Then, tissue homogenates or plasma were reacted with 200 μM pterin (Millipore Sigma) in black-walled 96-well plates (Greiner Bio-One) at 37°C for 60 min. The fluorescence was measured in a microplate reader (ARVO MX 1420; PerkinElmer) using excitation at 345 nm and emission detection at 390 nm. XO Activity was expressed as units/mg protein using native XO from buttermilk (Merck Millipore) as standard.

### Assessment of *in situ* ROS generation

We evaluated *in situ* generation of ROS using DHE, as previously described ^58^. The ascending aorta of human patients and mice was collected, and immediately embedded in Tissue-Tek OCT compound (Miles Laboratories, Inc.). DHE (5μM, FUJIFILM Wako Pure Chemical Corporation) was topically applied to the freshly frozen sections (5 μm), and incubated at 37°C for 30 min. The sections were stained and mounted in ProLong Gold Antifade Reagent with DAPI (Thermo Fisher Scientific). An all-in-one fluorescence microscope (BZ-9000; Keyence) was used to detect ROS generation as red fluorescence and elastic fibers as green autofluorescence.

### Assessment of *in situ* MMP activity

We measured gelatinase (MMP-2/gelatinase-A and MMP-9/gelatinase-B) activity by zymography. Fluorescein-conjugated DQ Gelatin from Pig Skin (50 μg/ml, Thermo Fisher Scientific) in reaction buffer (0.5 M Tris-HCl, pH 7.6, 1.5 M NaCl, 50 mM CaCl_2_, 2 mM NaN_3_) was applied to freshly frozen sections (5 μm), and incubated at 37°C for 24 h. Sections for negative control were incubated in the presence of 5 mM EDTA. The sections were stained and mounted in ProLong Gold Antifade Reagent with DAPI (Thermo Fisher Scientific). An all-in-one fluorescence microscope (BZ-9000; Keyence) was used to detect gelatinase (MMP-2 and MMP-9) activity as green fluorescence.

### Cell culture

Clonetics HAECs were purchased from Lonza, and maintained on a 1% gelatin-coated tissue culture dishes in M199 medium (MP Biomedicals) supplemented with 15% fetal bovine serum, 2 mM L-glutamine, 50 mg/ml heparin, and 30 mg/ml EC growth factor (Becton Dickinson). HAECs between the 5th and 8th passage were used for the experiments. Before hypotonic or cyclic stretch stimulation, cells were starved in EBM- 2 basal medium (Lonza, #CC-3162) with 0.5% fetal bovine serum for 12 h. SB-203580 (1 μM, LC Laboratories) and PF-573228 (20μM, Selleck Chemicals) were used for pharmacological inhibition of p38 MAPK and FAK, respectively.

The hypotonic solution was prepared by adding 30% distilled water to a modified Krebs solution (pH 7.3) containing 132 mM NaCl, 5.9 mM KCl, 1.2 mM MgCl_2_, 1.5 mM CaCl_2_, 11.5 mM glucose, 11.5 mM Hepes ^59^. Cyclic stretch stimulation was conducted using a mechanical strain loading device (STB-1400-10, STREX Inc.), as previously described ^60^. HAECs were seeded into a Cellmatrix Type I-P collagen (FUJIFILM Wako Pure Chemical Corporation) -coated silicone strain chambers (10 cm^2^, STB-CH- 10, STREX Inc.). The polydimethylsiloxane membrane was uniaxially and uniformly stretched (1 Hz, 10% elongation), or maintained under the static condition. For motion control, one end of the chamber was fixed by immobile plates to allow motion similar to cyclic stretch), while the other was fixed around a template that prevent the chamber from stretching. All experiments were performed at 37°C in a CO_2_ incubator.

### siRNA-mediated gene knockdown

HAECs were transiently transfected with EGR1 siRNA (#s4538, Ambion Silencer Select siRNAs, Thermo Fisher Scientific) and negative control siRNA (#4390843, Ambion Silencer Select Negative Control No. 1 siRNAs, Thermo Fisher Scientific) using HiPerFect Transfection Reagent (QIAGEN), according to the manufacturer’s instructions. Twenty-four h after transfection, cells were starved under a serum-free condition for 12 h before stimulation with hypotonic solution or cyclic stretch.

### Statistical analysis

All data are expressed as mean ± SEM. For two-group comparisons, two-tailed Student’s *t*-test was used. For multiple comparisons, one-way ANOVA with Tukey’s multiple comparisons test was used to compare each group. Values of *P* < 0.05 were considered statistically significant.

## ACKNOWLEDGMENTS

We thank Drs. Akifumi Kushiyama (Meiji Pharmaceutical University), Yoshito Yamashiro (University of Tsukuba), and Kimio Satoh (Tohoku University) for technical advice, and Drs. Daishi Fujita, Ryo Inuzuka, and Yuki Taniguchi for providing clinical information of MFS patients who underwent aortic root surgery.

## AUTHOR CONTRIBUTIONS

H.A. and I.K. conceived and designed the project; Hiroki Y. performed most of the experiments and analyzed data, with contributions from Q.L., K.Y., and H.T.; K.N., M.A., Haruo Y., and M.O. provided human tissue samples and collected clinical data; Norifumi T., Norihiko T., and M.A.F. provided experimental materials; A.S-K., M.U., H.K., R.M., and A.S. helped analysis and interpretation of the data: Hiroki Y., H.A., and I.K. wrote the manuscript. All authors read and approved the manuscript.

