## Supplementary Figures and Table for "Aberrant mechanosensitive signaling underlies activation of vascular endothelial xanthine oxidoreductase that promotes aortic aneurysm formation in Marfan syndrome"

[1.] Department of Cardiovascular Medicine, Graduate School of Medicine, The University of Tokyo, Tokyo, Japan. [2.] Marfan Syndrome Center, The University of Tokyo Hospital, Tokyo, Japan. [3.] Laboratory of System Physiology, Department of Biomedical Engineering, Graduate School of Medicine, The University of Tokyo, Tokyo, Japan. [4.] Department of Cardiac and Thoracic Surgery, St. Marianna University School of Medicine, Kawasaki, Japan. [5.] Department of Cardiac and Thoracic Surgery, Graduate School of Medicine, The University of Tokyo, Tokyo, Japan. [6.] Division of Cardiology and Metabolism, Center for Molecular Medicine, Jichi Medical University, Shimotsuke, Japan. [7.] Division of Pulmonary and Critical Care, Department of Medicine, Anschutz Medical Campus, University of Colorado-Denver School of Medicine, Aurora, CO, USA

#### Table of Contents

**SUPPLEMENTARY FIGURES**

Supplementary Fig. 1. Thoracic aortic aneurysm formation in *Fbn1* $^{C1039G/+}$ mice.

Supplementary Fig. 2. Thoracic aortic aneurysm formation in MFS patients.

Supplementary Fig. 3. Endothelial cells-specific knockout of XOR in *Fbn1* $^{C1039G/+}$ mice.

Supplementary Fig. 4. Effects of febuxostat on body weight, blood pressure, pulse rate, and blood biochemistry.

Supplementary Fig. 5. Schematic model depicting the contribution of endothelial XOR to the vicious cycle linking aberrant mechanosensitive signaling to the aortic phenotype in MFS.

**SUPPLEMENTARY TABLE**

Supplementary Table. Characteristic of patients whose ascending aortic tissues were used in this study.

### Supplementary Figure 1.

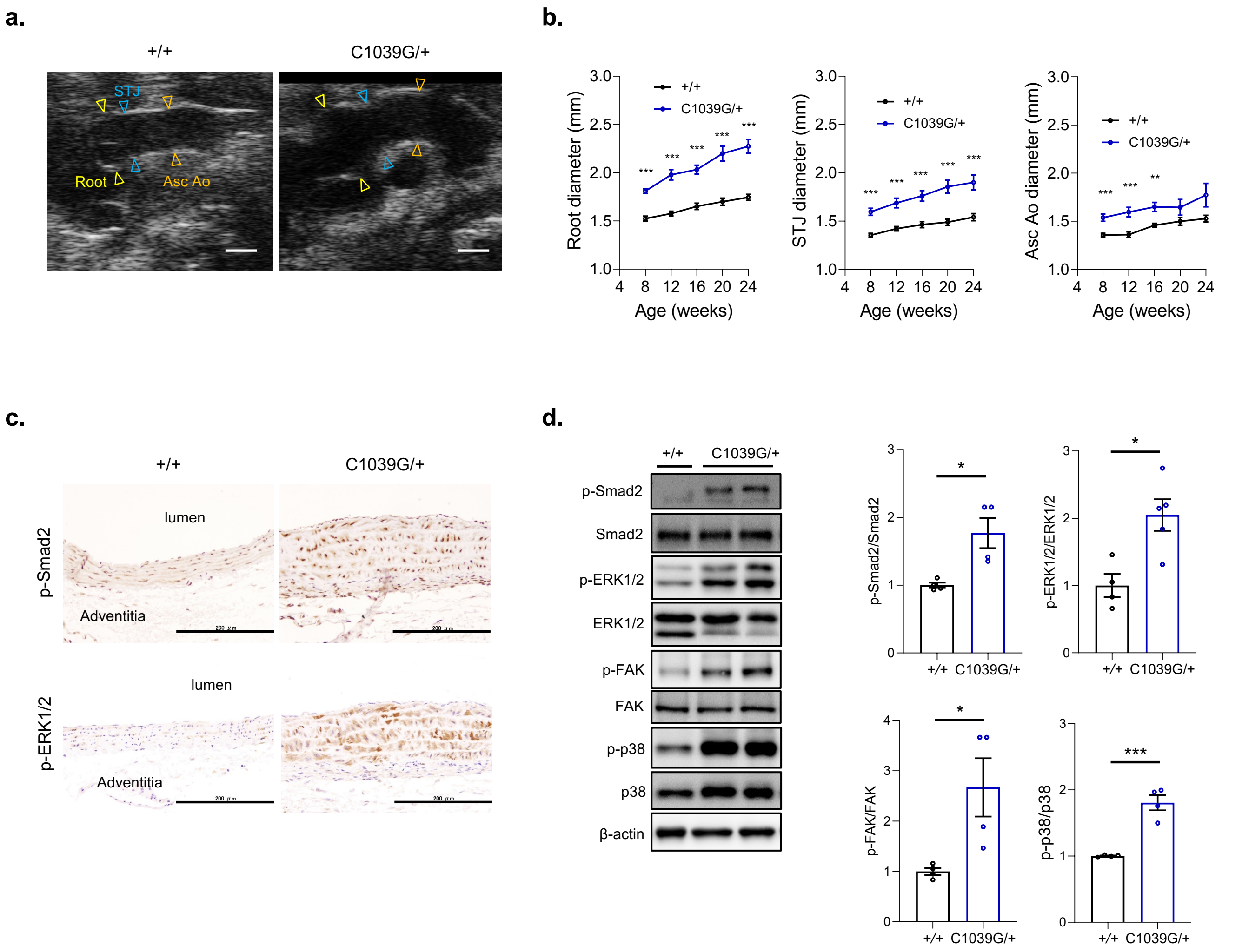


**Supplementary Figure 1. Thoracic aortic aneurysm formation in** ***Fbn1*** $^{C1039G/+}$ **mice.**
**a.** Ultrasound images of ascending aorta in *Fbn1*$^{+/+}$ and *Fbn1* $^{C1039G/+}$ mice (24 weeks of age). Arrowheads indicate aortic root (Root, yellow), sinotubular junction (STJ, blue), and ascending aorta (Asc Ao, orange). Scale bars, 1 mm.
**b.** Ascending aorta diameters at three different levels (Root, STJ, and Asc Ao) in *Fbn1*$^{+/+}$ (*n* = 10) and *Fbn1* $^{C1039G/+}$ mice (*n* = 11) (8 to 24 weeks of age). The data are presented as mean ± SEM. ****P*<0.001, two-tailed Student's *t*-test.
**c.** Immunostaining for phosphorylated Smad2 (p-Smad2) and phosphorylated ERK1/2 (p-ERK1/2) in ascending aorta of *Fbn1*$^{+/+}$ and *Fbn1* $^{C1039G/+}$ mice (24 weeks of age). Scale bars, 200 μm.
**d.** Immunoblot analysis of p-Smad2, Smad2, p-ERK1/2, ERK1/2, phosphorylated FAK (p-FAK), FAK, phosphorylated p38 MAPK (p-p38), p38 MAPK (p38), and β-actin in ascending aorta of *Fbn1*$^{+/+}$ and *Fbn1* $^{C1039G/+}$ mice. The intensity of each band was quantified by densitometric analysis, and quantitation graphs for the p-Smad2/Smad2 (*n* = 4), p-ERK1/2/ERK1/2 (*n* = 4 to 5), p-FAK/FAK (*n* = 4), and p-p38/p38 (*n* = 4) are shown (mean ± SEM). **P*<0.05, ****P*<0.001, two-tailed Student's *t*-test.

### Supplementary Figure 2.

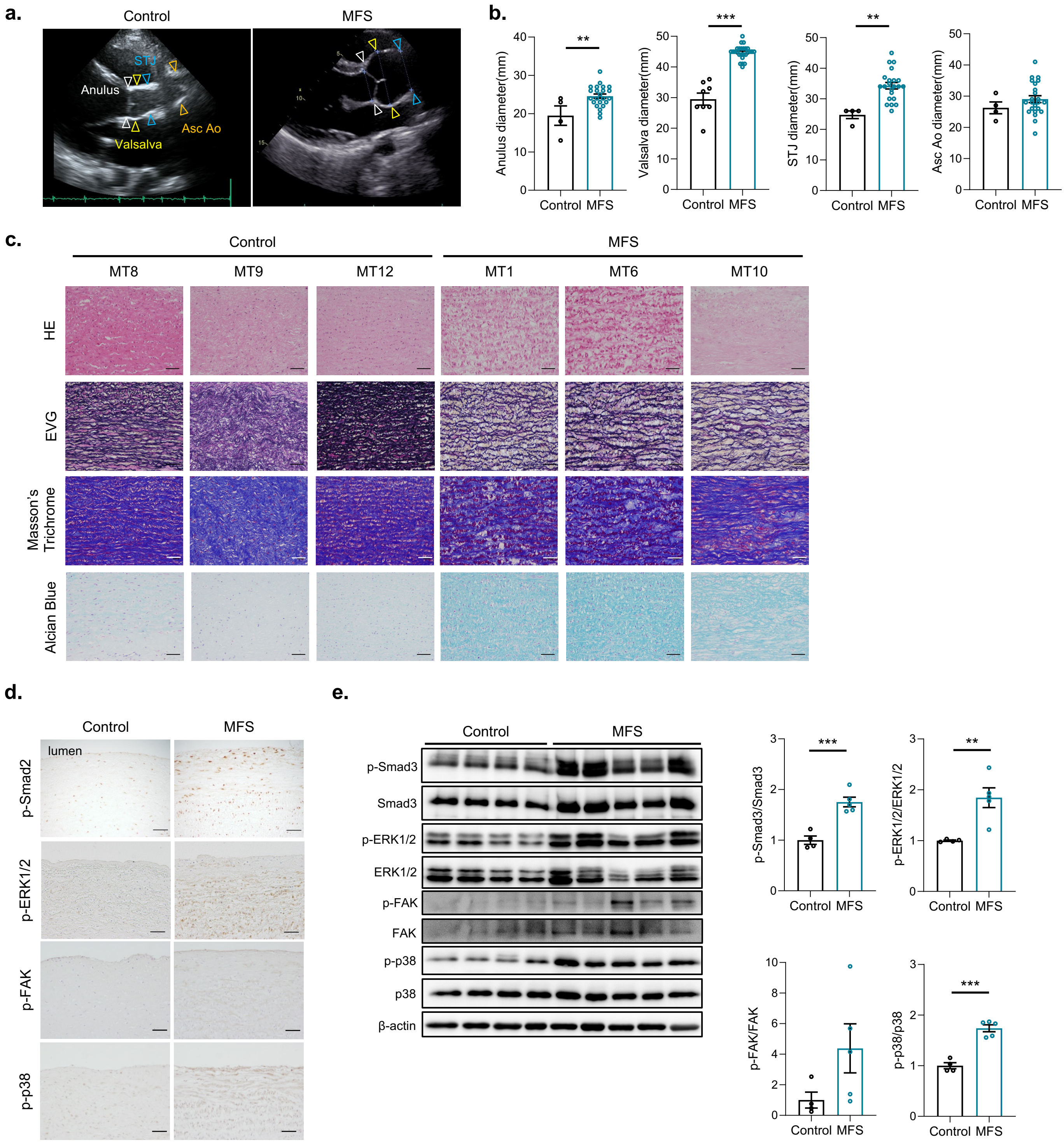


**Supplementary Figure 2. Thoracic aortic aneurysm formation in MFS patients.**

**a.** Ultrasound images of ascending aorta in MFS patients and control heart transplant recipients. Arrowheads indicate aortic annulus (Annulus, white), sinus of Valsalva (Valsalva, yellow), sinotubular junction (STJ, blue), and ascending aorta (Asc Ao, orange).

**b.** Ascending aorta diameters at four different levels (Annulus, Valsalva, STJ, and Asc Ao) in MFS patients ($n$ = 22 to 26) and control heart transplant recipients ($n$ = 4 to 8). The data are presented as mean ± SEM. $^{**}P<0.01$, $^{***}P<0.001$, two-tailed Student's *t*-test.

**c.** Histological analysis with HE staining, elastica van Gieson (EVG) staining, Masson's trichrome staining, and Alcian blue staining in ascending aorta of MFS patients (MT1, MT6, MT10, indicated in Supplemental Table) and control heart transplant recipients (MT8, MT9, MT12, indicated in Supplemental Table). Scale bars, 50 µm.

**d.** Immunostaining for phosphorylated Smad2 (p-Smad2), phosphorylated ERK1/2 (p-ERK1/2), phosphorylated FAK (p-FAK), and phosphorylated p38 MAPK (p-p38) in ascending aorta of MFS patient and control heart transplant recipient. Scale bars, 50 µm.

**e.** Immunoblot analysis of p-Smad3, Smad3, p-ERK1/2, ERK1/2, p-FAK, FAK, p-p38, p38, and β-actin in ascending aorta of MFS patients ($n$ = 5) and control heart transplant recipients ($n$ = 4). The intensity of each band was quantified by densitometric analysis, and quantitation graphs for the p-Smad2/Smad2, p-ERK1/2/ERK1/2, p-FAK/FAK, and p-p38/p38 are shown (mean ± SEM). $^{**}P<0.01$, $^{***}P<0.001$, two-tailed Student's *t*-test.

Supplementary Figure 3.

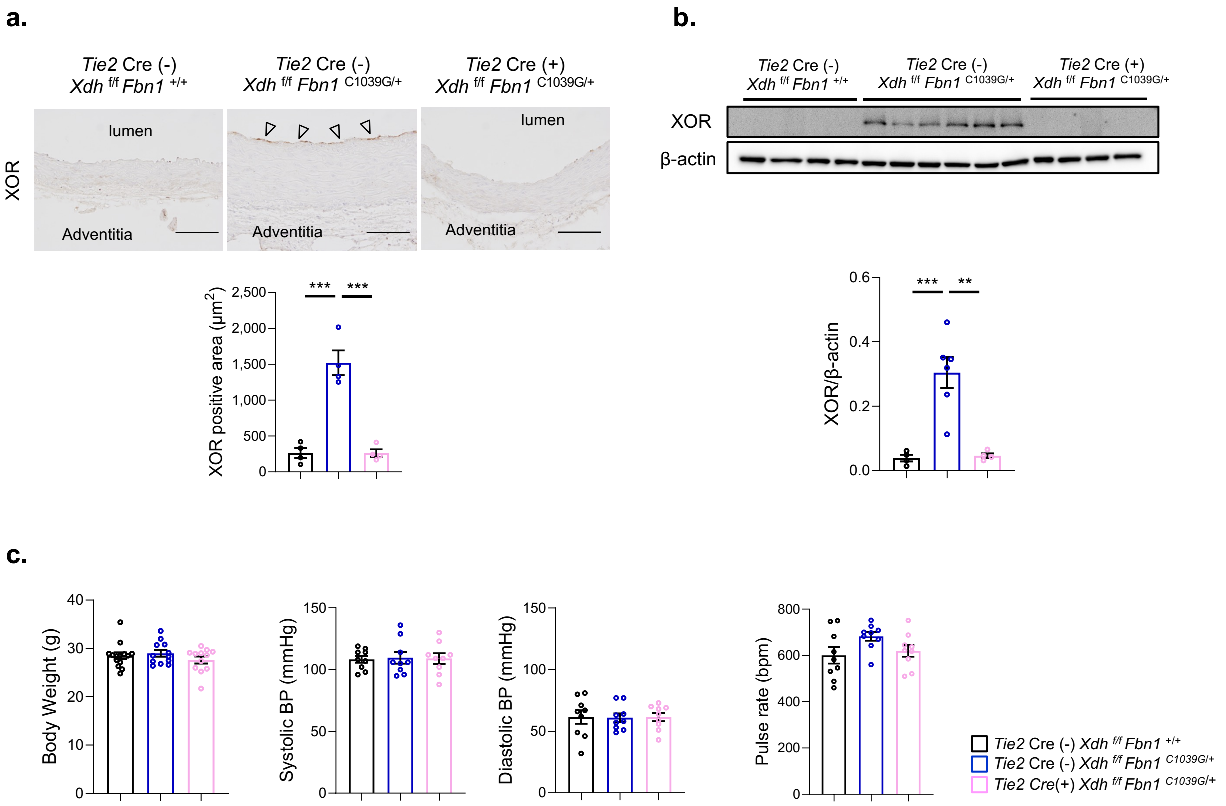


**Supplementary Figure 3. Endothelial cells-specific knockout of XOR in *Fbn1* $^{C1039G/+}$ mice.**
**a.** Immunostaining for XOR protein expression in ascending aorta of *Tie2* Cre (-) *Xdh* $^{f/f}$ *Fbn1* $^{+/+}$, *Tie2* Cre (-) *Xdh* $^{f/f}$ *Fbn1* $^{C1039G/+}$, and *Tie2* Cre (+) *Xdh* $^{f/f}$ *Fbn1* $^{C1039G/+}$ mice (16 weeks of age). Arrowheads indicate XOR expression in endothelial cells. Scale bars, 100 μm. The areas with XOR expression were quantified using an NIH Image J software (NIH, Research Branch), and quantitation graph (*n* = 4) is shown (mean ± SEM). ****P*<0.001, one-way ANOVA with Tukey's multiple comparisons test.
**b.** Immunoblot analysis of XOR expression in ascending aorta of *Tie2* Cre (-) *Xdh* $^{f/f}$ *Fbn1* $^{+/+}$ (*n* = 4), *Tie2* Cre (-) *Xdh* $^{f/f}$ *Fbn1* $^{C1039G/+}$ (*n* = 6), and *Tie2* Cre (+) *Xdh* $^{f/f}$ *Fbn1* $^{C1039G/+}$ mice (*n* = 4) (16 weeks of age). The intensity of each band was quantified by densitometric analysis and corrected for the amount of β-actin protein as an internal control (mean ± SEM). **P*<0.01, ***P*<0.001, one-way ANOVA with Tukey's multiple comparisons test.
**c.** Body weight (*n* = 12 to 14), blood pressures (BPs) (systolic, diastolic) (*n* = 9), and pulse rate (*n* = 9) of *Tie2* Cre (-) *Xdh* $^{f/f}$ *Fbn1* $^{+/+}$, *Tie2* Cre (-) *Xdh* $^{f/f}$ *Fbn1* $^{C1039G/+}$, and *Tie2* Cre (+) *Xdh* $^{f/f}$ *Fbn1* $^{C1039G/+}$ mice (16 weeks of age). The data are presented as mean ± SEM. **P*<0.01, ***P*<0.001, one-way ANOVA with Tukey's multiple comparisons test.

### Supplementary Figure 4.

**a.**

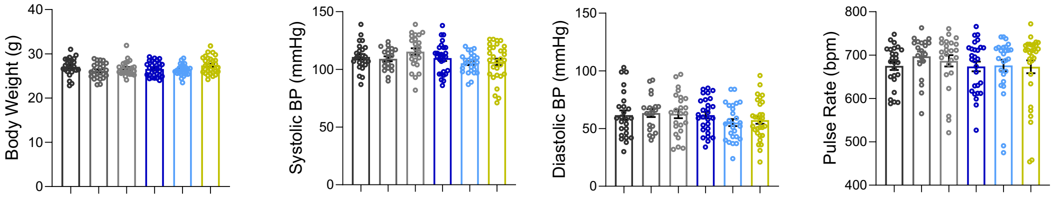


**b.**

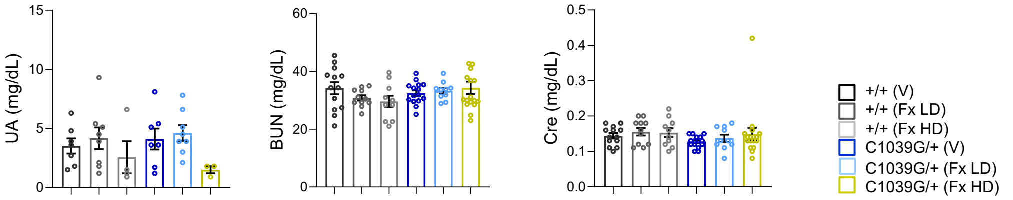


**Supplementary Figure 4. Effects of febuxostat on body weight, blood pressure, pulse rate, and blood biochemistry.**
**a.** Body weight (*n* = 24 to 30), blood pressures (BPs) (systolic, diastolic) (*n* = 20 to 30), and pulse rate (*n* = 20 to 30) of *Fbn1* $^{+/+}$ and *Fbn1* $^{C1039G/+}$ (16 weeks of age) after 8 weeks of treatment with vehicle (V), febuxostat at a lower dose of 1 mg/kg/day (Fx LD), or febuxostat at a higher dose of 5 mg/kg/day (Fx HD). The data are presented as mean ± SEM. One-way ANOVA with Tukey's multiple comparisons test.
**b.** Serum uric acid (UA; *n* = 4 to 8) and markers for kidney function（blood urea nitrogen BUN; *n* = 10 to 17, creatinine Cr; *n* = 10 to 17）in *Fbn1* $^{+/+}$ and *Fbn1* $^{C1039G/+}$ (16 weeks of age) after 8 weeks of treatment with V, Fx LD, or Fx HD.

Supplementary Figure 5.

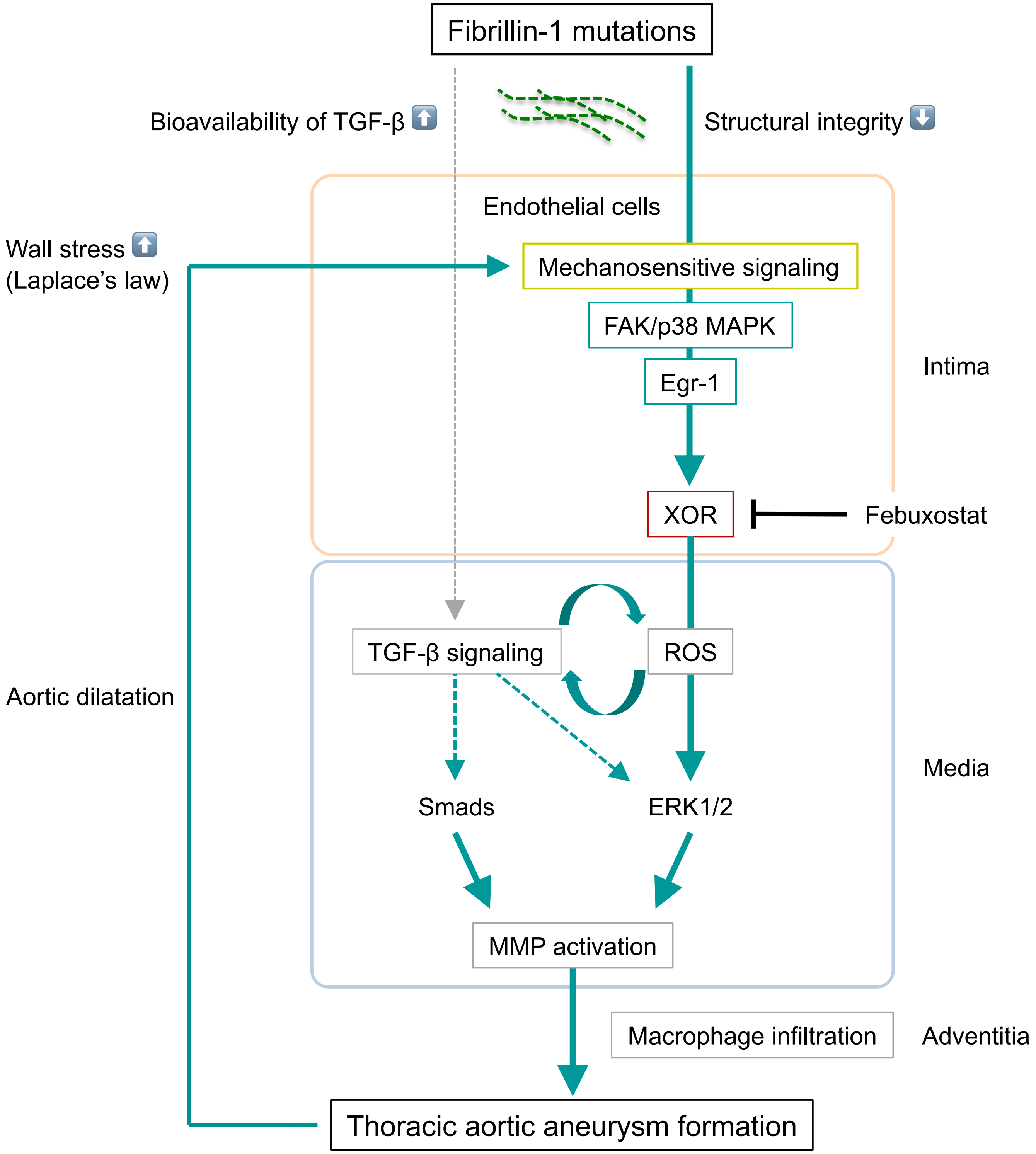


**Supplementary Figure 5. Schematic model depicting the contribution of endothelial XOR to the vicious cycle linking aberrant mechanosensitive signaling to the aortic phenotype in MFS.**
Mutations of Fibrillin-1, a major component of microfibrils in extracellular matrix, may alter the structural integrity of vascular wall, and aberrantly activate the mechanosensitive focal adhesion kinase (FAK)- p38 mitogen-activated protein kinase (MAPK) signaling pathway and up-regulate early growth response-1 (Egr-1), a mechanosensitive transcription factor, leading to up-regulation and activation of xanthine oxidoreductase (XOR) in endothelial cells. Endothelial XOR generates reactive oxygen species (ROS), which may amplify ROS generation by propagating signals in medial layer of aorta. Fibrillin-1 mutations also increase bioavailability of TGF-β, and ROS modulate TGF-β signaling in reciprocal manner in aortic wall. These pathological alterations induce matrix metalloproteinase (MMP) activation and macrophage infiltration, and thereby promote thoracic aortic aneurysm formation. An increase in the inner radius of aorta proportionally increases circumferential wall stress, and further activate mechanosensitive signaling in endothelial cells. A uric acid lowering drug febuxostat suppresses aortic aneurysm formation in Marfan syndrome (MFS) by inhibiting activation of endothelial XOR and thereby breaking the vicious cycle linking aberrant mechanosensitive signaling to the aortic phenotype. ERK; extracellular signal-regulated kinase.

**Supplementary Table. Characteristic of patients whose ascending aortic tissues were used in this study.**

| MFS patient | Age | Sex | Aortic diameter (mm) (Anulus/Valsalva/STJ/Asc aorta) | Nucleotide change | Predicted protein change |
|---|---|---|---|---|---|
| MT1 | 42 | F | 22 / 41 / 30 / 31 | c.6268G>T | p.Glu2090Ter |
| MT5 | 54 | F | 21 / 45 / 45 / 35 | c.2237A>G | p.Tyr746Cys |
| MT6 | 45 | F | 20 / 45 / 28 / 26 | c.5368C>T | p.Arg1790Ter |
| MT7 | 21 | M | 27 / 46 / 36 / 28 | c.6740-2A>G | Exon 56 deletion |
| MT10 | 57 | M | 24 / 45 / 34 / 36 | c.1786T>G | p.Cys596Gly |
| MT20 | 10 | F | 26 / 44 / 36 / 18 | c.3458G>A | p.Cys1153Tyr |
| MT25 | 49 | M | — / 50 / — / — | — | — |
| MT27 | 23 | F | — / 41 / 35 / 25 | n.d. | n.d. |
| MT28 | 26 | M | 28 / 45 / 34 / 22 | c.8326C>T | p.Arg2776Ter |
| MT29 | 31 | M | 19 / 45 / 36 / 29 | — | — |
| MT31 | 27 | F | 26 / 44 / 28 / 25 | c.3544T>G | p.Cys1182Gly |
| MT32 | 34 | M | 27 / 48 / 42 / 41 | — | — |
| MT33 | 36 | F | — / 41 / — / — | c.3770T>A | p.Ile1257Asn |
| MT34 | 66 | F | 23 / 45 / 32 / 29 | c.4621C>T | p.Arg1541Ter |
| MT35 | 19 | M | 24 / 44 / 30 / 25 | c.7327_7330+2delGTAGGT | Exon 59 deletion |
| MT36 | 17 | M | 24 / 46 / 37 / 35 | c.3919T>C | p.Cys1307Arg |
| MT38 | 28 | F | 27 / 46 / 33 / 28 | — | — |
| MT41 | 22 | M | 23 / 43 / 34 / 24 | n.d. | n.d. |
| MT42 | 20 | F | 22 / 40 / — / — | c.5788+5G>C | Exon 47 deletion |

| | | | | | |
|---|---|---|---|---|---|
| MT43 | 22 | M | 23 / 45 / 42 / 28 | c.7565delG | p.Cys2522Serfs*160 |
| MT44 | 34 | M | 26 / 45 / — / 32 | n.d. | n.d. |
| MT45 | 27 | M | 24 / 48 / 26 / 28 | c.239G>A | p.Cys80Tyr |
| MT46 | 41 | F | 26 / 46 / 40 / 36 | n.d. | n.d. |
| MT47 | 18 | M | 31 / 46 / 34 / 27 | c.2033_2066delTTTG | p.V675fsTer |
| MT48 | 24 | M | 25 / 44 / 28 / 26 | c.3524_3525delTA | p.Ile1175fs |
| MT49 | 33 | F | 27 / 46 / 34 / 34 | c.2645C>T | p.Ala882Val |

| Control patient | Age | Sex | Aortic diameter (mm) (Anulus/Valsalva/STJ/Asc aorta) | Underlying cardiac disease |
|---|---|---|---|---|
| MT8 | 59 | F | — / 35 / — / — | DCM |
| MT9 | 43 | M | 18 / 32 / 26 / 31 | DCM |
| MT11 | 45 | M | — / 33 / — / — | ICM |
| MT12 | 33 | M | 13 / 19 / 16 / 22 | DCM |
| MT21 | 43 | M | 23 / 27 / 21 / 25 | DCM |
| MT22 | 31 | M | — / 27 / — / — | DCM |
| MT23 | 53 | M | 24 / 36 /26 / 27 | DCM |
| MT24 | 18 | M | — / 27 / — / — | DCM |

Asc aorta, ascending aorta; MFS, Marfan syndrome; M, male; F, female; DCM, dilated cardiomyopathy; ICM, ischemic cardiomyopathy; STJ, sinotubular junction.
